## Supplemental Tables and Figures for "Functional assessment of the “two-hit” model for neurodevelopmental defects in *Drosophila* and *X. laevis*": S1 Table.pdf

| HGNC symbol | Organism | Homolog | % identity | DIOPT score | DIOPT rank | Larval central nervous system expression (FlyAtlas) |
|---|---|---|---|---|---|---|
| *UQCRC2* | *Drosophila* | *UQCR-C2* | 31% | 13.82 | High | Moderate |
| *CDR2* | *Drosophila* | *Cen* | 12% | 8.93 | High | Moderate |
| *MOSMO* | *Drosophila* | *CG14182* | 51% | 11.89 | High | Low |
| *POLR3E* | *Drosophila* | *Sin* | 32% | 12.83 | High | Moderate |
| *EEF2K* | *Drosophila* | NA | NA | NA | NA | NA |
| *VWA3A* | *Drosophila* | NA | NA | NA | NA | NA |
| *PDZD9* | *Drosophila* | NA | NA | NA | NA | NA |
| *UQCRC2* | *X. laevis* | *uqcrc2* | 70% | --- | --- | --- |
| *CDR2* | *X. laevis* | *cdr2* | 63% | --- | --- | --- |
| *MOSMO* | *X. laevis* | *mosmo* | 82% | --- | --- | --- |
| *POLR3E* | *X. laevis* | *polr3e* | 66% | --- | --- | --- |
| *EEF2K* | *X. laevis* | *eef2k* | 72% | --- | --- | --- |
| *VWA3A* | *X. laevis* | *vwa3a* | 51% | --- | --- | --- |
| *PDZD9* | *X. laevis* | NA | NA | --- | --- | --- |
