## Supplemental Tables and Figures for "Functional assessment of the “two-hit” model for neurodevelopmental defects in *Drosophila* and *X. laevis*": S2 Table_revision.pdf

### Table S2A

| Genes modulating phenotypes of 16p12.1 homologs[#] | Total | 16p12.1 homologs | Patient-specific "second-hit" genes | Neurodevel/ functionally related genes | Transcriptome targets |
|---|---|---|---|---|---|
| Total number of crosses | 521 | 39 | 227 | 119 | 136 |
| RNAi/mutant/overexp lines | 134 | 10 | 46 | 24 | 54 |
| Genes (fly homologs) | 80 | 4 | 24 | 13 | 39 |
| Tested pairwise combinations of RNAi lines | 388 | 30 | 181 | 95 | 82 |
| Tested pairwise combinations of genes | 224 | 12 | 96 | 55 | 61 |
| Pairwise combinations enhancing phenotypes due to knockdown of 16p12.1 homologs | 54 | 3 | 26 | 6 | 19 |
| Pairwise combinations suppressing phenotypes due to knockdown of 16p12.1 homologs | 29 | 1 | 6 | 12 | 10 |
| Pairwise combinations that do not affect the phenotype of 16p12.1 homologs | 76 | 3 | 32 | 17 | 24 |
| Pairwise combinations whose effects on 16p12.1 homologs were not validated | 65 | 5 | 32 | 20 | 8 |

### Table S2B

| Interactions* | Total | 16p12.1 homologs | Patient-specific "second-hit" genes | Neurodevel/ functionally related genes | Transcriptome targets |
|---|---|---|---|---|---|
| Negative interactions (confirmed and potential) | 41 | 0 | 6 | 10 | 25 |
| Positive interactions (confirmed and potential) | 63 | 3 | 31 | 12 | 17 |
| No interaction (confirmed and potential) | 61 | 2 | 30 | 18 | 11 |
| Not validated interaction | 55 | 7 | 29 | 12 | 7 |

[#]Genes modulating phenotypes of 16p12.1 homologs were confirmed using Mann-Whitney tests.
*Genetic interactions identified using the multiplicative model. Genes with no single knockdown *Flynotyper* phenotypes were excluded from multiplicative model testing.
