## Supplemental Tables and Figures for "Functional assessment of the “two-hit” model for neurodevelopmental defects in *Drosophila* and *X. laevis*": S3 Table_revision.pdf

| **EXPERIMENT** | **Individual knockdown of 16p12.1 homologs** | **Pairwise knockdown of 16p12.1 homologs** | **Pairwise interactions of 16p12.1 homologs with "Neurodevelopmental genes"** | **Pairwise interactions of 16p12.1 homologs with "Transcriptome targets"** | **Pairwise interactions of 16p12.1 homologs with "Patient-specific second-hit genes"** | **Cellular phenotypes** |
|---|---|---|---|---|---|---|
| **HYPOTHESIS TESTED** | Multiple 16p12.1 homologs contribute to neurodevelopmental phenotypes | 16p12.1 homologs interact towards neurodevelopmental phenotypes | 16p12.1 homologs interact with genes within core neurodevelopmental pathways and functionally related genes towards neurodevelopmental phenotypes | 16p12.1 homologs interact with their downstream genes towards neurodevelopmental phenotypes | Patient-specific “second-hits” can modulate neurodevelopmental phenotypes of 16p12.1 homologs through genetic interactions | Genes interact with 16p12.1 homologs towards neuronal defects by altering cellular phenotypes |
| **EXPERIMENTAL STRATEGY** | Individual knockdown of 16p12.1 homologs using tissue-specific drivers in *Drosophila* and whole-embryo knockdown in *X. laevis*, and assessment of multiple neuronal, cellular and developmental phenotypes | Simultaneous knockdown of 16p12.1 homologs - assessment of genetic interactions in the fly eye. Assessment of specific pairs of homologs towards brain and cellular phenotypes in *X. laevis* | Simultaneous knockdown of 16p12.1 homolog with neurodevelopmental genes and functionally related genes - assessment for genetic interactions in the fly eye | Simultaneous knockdown of 16p12.1 homolog with transcriptional targets or their functionally related genes - assessment for genetic interactions in the fly eye | Simultaneous knockdown of 16p12.1 homologs with homologs of patient-specific “second-hit” genes - assessment for genetic interactions in the fly eye | Assessment of cellular proliferation and apoptosis processes for validated two-hit interactions in the developing fly eye |
| | | | Genes in established neurodevelopmental pathways - 7 genes; Genes functionally related to 16p12.1 homologs - 6 genes | Up and down-regulated genes from transcriptome analysis of 16p12.1 homologs - 25 genes; Genes within enriched GO terms from transcriptome analysis - 14 genes | Homologs of “second-hit” genes from sequencing children with 16p12.1 deletion - 24 genes | |
| | | 12 combinations | 55 combinations | 61 combinations | 96 combinations | 3 combinations |
| **OUTCOME** | Global and homolog-specific phenotypes | 3/12 positive interactions | 22/55 interactions | 42/61 interactions | 37/96 interactions | Alterations of proliferation and apoptosis processes compared to single knockdown of 16p12.1 homolog |

| EXPERIMENT | Individual knockdown of 16p12.1 homologs | Pairwise knockdown of 16p12.1 homologs | Pairwise interactions of 16p12.1 homologs with "Neurodevelopmental genes" | Pairwise interactions of 16p12.1 homologs with "Transcriptome targets" | Pairwise interactions of 16p12.1 homologs with "Patient-specific second-hit genes" | Cellular phenotypes |
|---|---|---|---|---|---|---|
| **CONCLUSIONS** | Each 16p12.1 homolog contributes towards a range of developmental, neuronal, and cellular defects in flies/*X. laevis* | 16p12.1 homologs contribute towards neurodevelopmental phenotypes through mild genetic interactions and additive effects | 16p12.1 homologs interact with genes in core pathways towards neurodevelopmental phenotypes | Differentially expressed genes interact with 16p12.1 homologs towards neurodevelopmental phenotypes | Patient-specific second-hits can modify phenotypes of 16p12.1 homologs through synergistic and alleviating interactions | Interacting genes modulate proliferation and apoptosis phenotypes of 16p12.1 homologs during development |
| **HYPOTHESIS-GENERATING RESULTS** | Validation of phenotypes associated with each homologs in other model organisms | Assessment of identified interactions in specific neuronal cell types | Follow up of identified interactions (such as those involving homologs of *MOSMO* and *SETD5*) in other model organisms and human cell lines/organoids. | | | Assessment of mechanisms connecting genes and interactions to cellular defects in specific neuronal cell types |
