## Supplemental Tables and Figures for "Functional assessment of the “two-hit” model for neurodevelopmental defects in *Drosophila* and *X. laevis*": S4 Figure_revision.pdf

**A**

| | Immune system process | Response to stimulus | Cuticle developemnt | Protein folding | Heat response | Metabolic process | Muscle contraction | Response to oxygen | Cell adhesion | Drug metabolism | Pigmentation | Respiratory system dev. |
|---|---|---|---|---|---|---|---|---|---|---|---|---|
| *Cen* $^{GD9689}$ | ■ | ■ | ■ | ■ | ■ | | ■ | ■ | | | | |
| *CG14182* $^{GD2738}$ | ■ | ■ | | | | ■ | | | | | | |
| *Sin* $^{GD7027}$ | | | ■ | | | ■ | | | ■ | ■ | ■ | ■ |
| *UQCR-C2* $^{GD11238}$ | ■ | ■ | | ■ | ■ | | | | | | | |

| | Cellular respiration | Metabolic process | Proteolysis | Cell adhesion | Circulatory process | Drug metabolism | Immune system process | Response to stimulus | Signaling regulation | Synaptic assembly | Synaptic transmission | Vesicle transport | Cell differentiation | Homeostasis | Molecular transport | Muscle contraction | Nervous system dev. | Neuron proliferation | Sensory perception | Spindle organization | System/organ dev. | Protein folding |
|---|---|---|---|---|---|---|---|---|---|---|---|---|---|---|---|---|---|---|---|---|---|---|
| *Cen* $^{GD9689}$ | ■ | ■ | ■ | | | | | | | | | | | | | | | | | | | |
| *CG14182* $^{GD2738}$ | ■ | ■ | ■ | ■ | ■ | ■ | ■ | ■ | ■ | ■ | ■ | ■ | | | | | | | | | | |
| *Sin* $^{GD7027}$ | ■ | ■ | ■ | ■ | ■ | ■ | ■ | ■ | | | | | ■ | ■ | ■ | ■ | ■ | ■ | ■ | ■ | ■ | |
| *UQCR-C2* $^{GD11238}$ | ■ | ■ | | | | | | | | | | | | | | | | | | | | ■ |

**B**

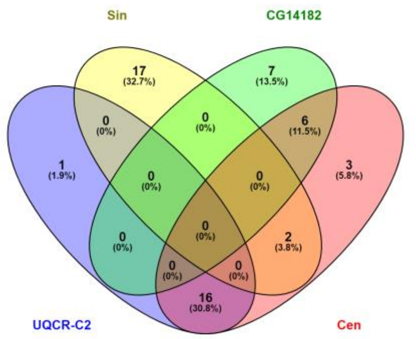


**C**

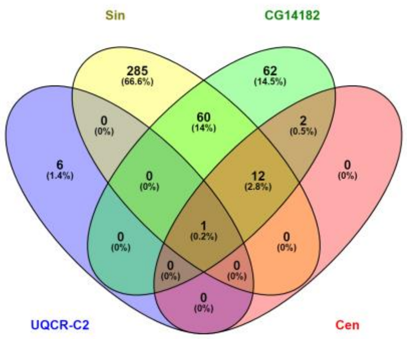


**D**

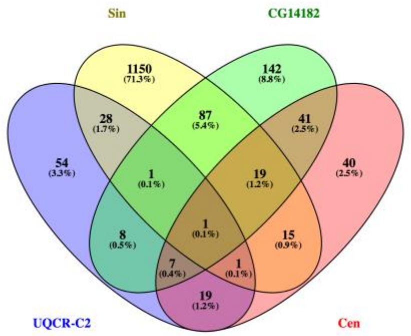


**E**

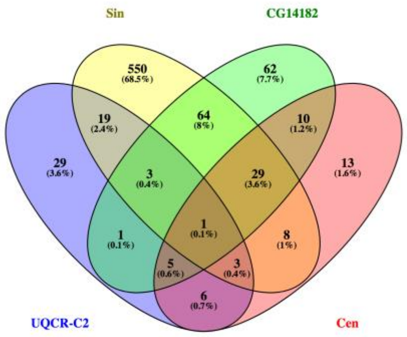
