## Supplemental Tables and Figures for "Functional assessment of the “two-hit” model for neurodevelopmental defects in *Drosophila* and *X. laevis*": S4 Table_revision.pdf

| Human gene | Chr. | Position | Ref. | Alt. | Variant Type | Family | Phenotype change of 16p12.1 homologs | Genetic interaction with 16p12.1 genes |
|---|---|---|---|---|---|---|---|---|
| *SETD5* | chr3 | 9488832 | T | TAC | Frameshift insertion | GL_01 | Enhancer *Sin*; Enhancer *CG14182* | Positive: *Cen* Negative: *CG14182* |
| *LAMC3* | chr9 | 133927967 | C | T | Stopgain | GL_01 | Enhancer *UQCR-C2* | Positive: *UQCR-C2* and *Sin* |
| *DMD* | chrX | 33229421 | C | T | Stopgain | GL_01 | No change/Not validated | Positive: *Sin* |
| *DPM1* | chr20 | 49574926 | C | T | Stopgain | GL_01 | No change/Not validated | No interaction/Not validated |
| *DNAH7* | chr2 | 196759808 | A | AT | Stopgain | GL_07 | No change/Not validated | No interaction/Not validated |
| *PDE11A* | chr2 | 178879181 | G | A | Stopgain | GL_07 | No change/Not validated | No interaction/Not validated |
| *CEP135* | chr4 | 56877651 | CAG | C | Frameshift deletion | GL_11 | Suppressor *UQCR-C2* | Negative: *Cen* and *Sin* |
| *NRXN1* | chr2 | 50837494 | NA | NA | Deletion | GL_11 | Enhancer *UQCR-C2* and *Sin* | Positive: *CG14182* |
| *PDLIM5* | chr4 | 95575739 | A | G | Nonsynonymous | GL_12 | Enhancer *Cen*, *CG14182* | Positive: *UQCR-C2* and *Sin* |
| *ARID1B* | chr6 | 157099425 | A | AGC | Frameshift insertion | GL_13 | Suppressor *UQCR-C2*, *Cen*, *CG14182*, and *Sin* | Positive: *Cen* and *CG14182* |
| *PEX1* | chr7 | 92123885 | GAGGAGCA | G | Frameshift deletion | GL_13 | Enhancer *Sin* | Positive: *CG14182* |
| *PYGM* | chr11 | 64527223 | G | A | Stopgain | GL_15 | No change/Not validated | Potential positive *UQCR-C2*, *Cen*, *Sin*, *CG14182* |
| *DST* | chr6 | 56480833 | G | A | Stopgain | GL_19 | Enhancer *UQCR*-C2, *Cen*, *Sin*, and *CG14182* | Positive: *UQCR-C2*, *Cen*, *Sin* |
| *PSMD1* | chr2 | 231949783 | T | A | Stopgain | GL_19 | Enhancer *UQCR-C2*, *Sin*, and *CG14182* | Potential positive: *UQCR-C2*, *Sin* |
| *RAPGEF6* | chr5 | 130771676 | G | A | Stopgain | GL_20 | Enhancer *UQCR-C2*, *Cen*, *Sin*, and *CG14182* | Positive: *UQCR-C2*, *CG14182* |
| *NALCN* | chr13 | 101763037 | G | C | Stopgain | GL_22 | No change/Not validated | No interaction/Not validated |
| *CADPS* | chr3 | 62739381 | C | T | Nonsynonymous | GL_22 | Enhancer *Sin* | Positive: *UQCR-C2* and *Sin* |
| *CAPN9* | chr1 | 230895256 | A | G | Splicing | GL_33 | Enhancer *UQCR-C2* and *CG14182* | Potential positive: *Sin* |

| Human gene | Chr. | Position | Ref. | Alt. | Variant Type | Family | Phenotype change of 16p12.1 homologs | Genetic interaction with 16p12.1 genes |
|---|---|---|---|---|---|---|---|---|
| *CHRNA7* | chr15 | 30936285 | NA | NA | Deletion | GL_36 | Enhancer *Cen* | Potential negative *Cen*, *Sin*, *CG14182* |
| *USP45* | chr6 | 99891524 | C | A | Stopgain | GL_46 | No change/Not validated | Potential no interaction |
| *PDE11A* | chr2 | 178681632 | CA | C | Frameshift deletion | GL_46 | No change/Not validated | No interaction/Not validated |
| *DNAH10* | chr12 | 124289588 | G | C | Splicing | GL_48 | Enhancer *UQCR-C2* and *Sin*, Enhancer *CG14182* | Potential positive: *UQCR-C2*, *Cen*, *CG14182* |
| *CACNA1A* | chr19 | 13342531 | G | A | Nonsynonymous | GL_51 | Suppressor *CG14182* | Potential positive *UQCR-C2, Cen*, *Sin*, *CG14182* |
| *OPRL1* | chr20 | 62729293 | GAC | G | Frameshift deletion | GL_52 | No change/Not validated | No interaction/Not validated |
