## Supplemental Tables and Figures for "Functional assessment of the “two-hit” model for neurodevelopmental defects in *Drosophila* and *X. laevis*": S5 Table.pdf

| Experiment | | Knockdown of *Drosophila* homologs of 16p12.1 genes | | | |
|---|---|---|---|---|---|
| **Phenotype** | **Assay** | *UQCR-C2* | *Cen* | *Sin* | *CG14182* |
| Adult eye morphology | Eye phenotype (overexpression of *Dicer2*) | Normal | Normal | Rough eye | Rough eye |
| | Eye phenotype (no overexpression of *Dicer2*) | Normal | Normal | Moderate rough eye | Normal |
| Role in development | Ubiquitous knockdown | Larval lethal | Normal | Larval lethal | Normal |
| | Wing development | Lethal | Normal | Severe phenotype | Normal |
| Neuronal phenotypes | Lifespan | Increased | Normal | Reduced | Reduced |
| | Developmental timing | Normal | Normal | Delayed / larval lethality | Partial larval lethality |
| | Seizure susceptibility | Increased | Normal | Normal | Normal |
| | Complexity of dendritic arbors | Normal | Normal | Normal | Reduced |
| | Brain size | Normal | Normal | Reduced | Reduced |
| Cellular proliferation (developing brain) | pH3 staining | NA | NA | Reduced | Reduced |
| Apoptosis (developing brain) | Dcp-1 staining | NA | NA | Reduced | Normal |
| RNA sequencing (adult heads) | Differential gene expression (fly homologs) | Protein folding, heat shock | Protein folding, heat shock protein, muscle contraction | Cell adhesion, respiratory system development | No clear relevant functional enrichment |
| | Differential gene expression (human homologs) | Protein folding | Proteolysis | Muscle contraction, nervous system development, system/organ development | Synapse assembly and transmission, histone methyltransferase function, small nucleolar ribonuclear complex |

| Experiment | | Knockdown of *X. laevis* homologs of 16p12.1 genes | | | |
|---|---|---|---|---|---|
| **Phenotype** | **Assay** | *uqcrc2* | *cdr2* | *polr3e* | *mosmo* |
| Craniofacial features | Face width | Normal | Decreased | Decreased | Decreased |
| | Face height | Normal | Increased | Normal | Increased |
| | Orofacial area | Normal | Normal | Decreased | Decreased |
| | Eye area | Normal | Decreased | Decreased | Decreased |
| | Face angle | Normal | Decreased | Decreased | Decreased |
| Brain phenotypes | Forebrain size (partial KD) | Normal | Normal | Normal | Reduced |
| | Midbrain size (partial KD) | Normal | Normal | Normal | Reduced |
| | Forebrain size (stronger KD) | Normal | Lethal | Reduced | Reduced |
| | Midbrain size (stronger KD) | Normal | Lethal | Reduced | Reduced |
| Axon outgrowth | Axon length (stronger KD) | Normal | Lethal | Normal | Decreased |
| Cellular proliferation (developing embryo) | pH3 staining - western blot | NA | NA | Reduced | Normal |
