## Supplemental Tables and Figures for "Functional assessment of the “two-hit” model for neurodevelopmental defects in *Drosophila* and *X. laevis*": S6 Table.pdf

| Organism | Human gene | Gene | Primer name | Primer sequence (5'-3') |
|---|---|---|---|---|
| *D. melanogaster* | *UQCRC2* | *UQCR-C2* | UQCR-C2_Fwd1 | TCTGTCAAGGCTGTGAATGCC |
| *D. melanogaster* | | | UQCR-C2_Rev1 | AAAACCGAACAGACCAGCGT |
| *D. melanogaster* | *CDR2* | *Cen* | Cen_Fwd1 | GCAGACGGACAACTCCATCC |
| *D. melanogaster* | | | Cen_Rev1 | TCACCATGGGAGAGCCATTC |
| *D. melanogaster* | *MOSMO* | *CG14182* | CG14182_Fwd | TCCCGACTGGATCATCACG |
| *D. melanogaster* | | | CG14182_Rev | AGTCCCAATCGGATGTCACC |
| *D. melanogaster* | *POLR3E* | *Sin* | Sin_Fwd1 | AAACGTGGCATCATGGACAA |
| *D. melanogaster* | | | Sin_Rev1 | GGTTATGGAACGCGAGCTTG |
| *D. melanogaster* | *RP49* | *Rp49* | rp49_Fwd | GCAAGCCCAAGGGTATCGA |
| *D. melanogaster* | | | rp49_Rev | ACCGATGTTGGGCATCAGA |
| *X. laevis* | *MOSMO* | *mosmo* | MOSMO_Fwd_L_S | CTTTGCCATCGCCAGTATCG |
| *X. laevis* | | | MOSMO_Rev_L | GGTAATTTGTAGGGTTGGCCTC |
| *X. laevis* | | | MOSMO_Rev_S | GGATGTTTGTTTCTGGCAGC |
| *X. laevis* | *UQCRC2* | *uqcrc2* | UQCRC2_Fwd_L | CCGTGGAATTGAAGCTGTTG |
| *X. laevis* | | | UQCRC2_Rev_L | TAATCCAACCAGTGCCATCC |
| *X. laevis* | | | UQCRC2_Fwd_S | ATTACTCGCCCTCATCCAAG |
| *X. laevis* | | | UQCRC2_Rev_S | CAGTACAAGGAGTTAGCCAGTG |
| *X. laevis* | *POLR3E* | *polr3e* | POLR3E_Fwd_L | GGAAAATGAAGATGACGATCC |
| *X. laevis* | | | POLR3E_Rev_L | GATGTGCCATCAACATTCAGG |
| *X. laevis* | *CDR2* | *cdr2* | CDR2_Fwd_L_S | GACAGCAACGTGGAGGAGTTC |
| *X. laevis* | | | CDR2_Rev_L_S | TGCTCATTCATCCGACGCAG |
| *X. laevis* | *SETD5* | *setd5* | SETD5_Fwd_L_S | ATCCCTCTGGGAGTCACCAC |
| *X. laevis* | | | SETD5_Rev_L_S | TGAGTGAATCCCATTGTGCTCTG |
| *X. laevis* | *ODC1* | *ODC1* | ODC1_Fwd | GCCATTGTGAAGACTCTCTCCATTC |
| *X. laevis* | | | ODC1_Rev | TTCGGGTGATTCCTTGCCAC |
