## Supplemental Tables and Figures for "Functional assessment of the “two-hit” model for neurodevelopmental defects in *Drosophila* and *X. laevis*": S7 Table.pdf

| Morpholino target | Morpholino sequence |
|---|---|
| *mosmo* L allele | 5-ACAATTGACATCCACTTACTGCCGG-3 |
| *mosmo* S allele | 5- CACCTTCCCTACCCCGCTACTTAC-3 |
| *polr3e* L allele | 5-ACTGTAAGCCTCTTTTGCCTTACCT-3 |
| *uqcrc2* L allele | 5-ACAGTGTCTCTAAAGCACAGATACA-3 |
| *uqcrc2* S allele | 5-CCCCTAACCCATTAAACATATACCT-3 |
| *cdr2* S and L alleles | 5-CATCCCTCCCATACACTCACCTTG-3 |
| *setd5* S and L alleles | 5-TGATTCAGGGCTGTAGAGGAGAAAA-3 |
| standard control morpholino | 5-CCTCTTACCTCAGTTACAATTTATA-3 |
