## Supplementary figures and images for "Functional assessment of the “two-hit” model for neurodevelopmental defects in *Drosophila* and *X. laevis*"

### S1 Figure_revision.pdf

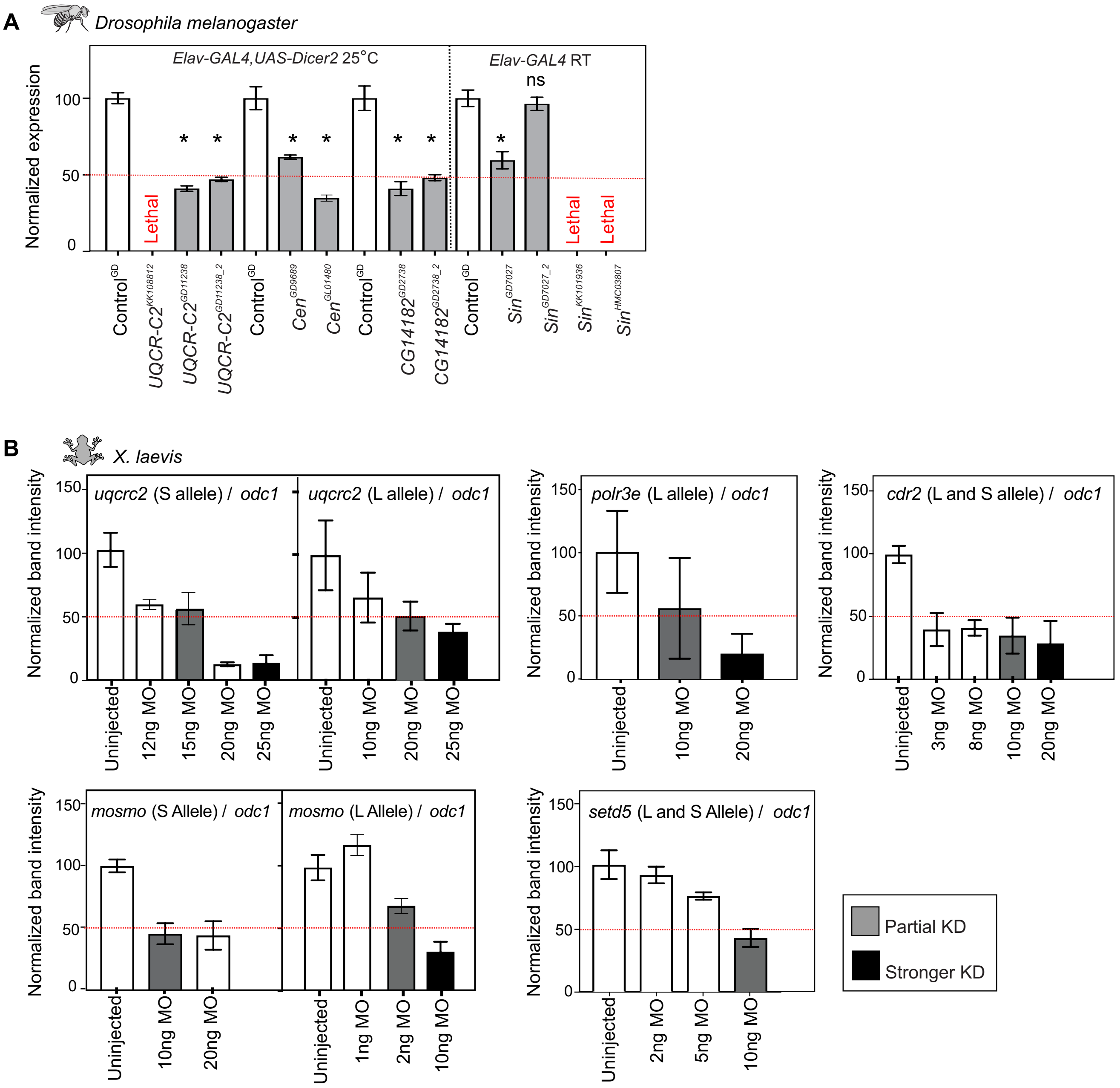

### S2 Figure.pdf

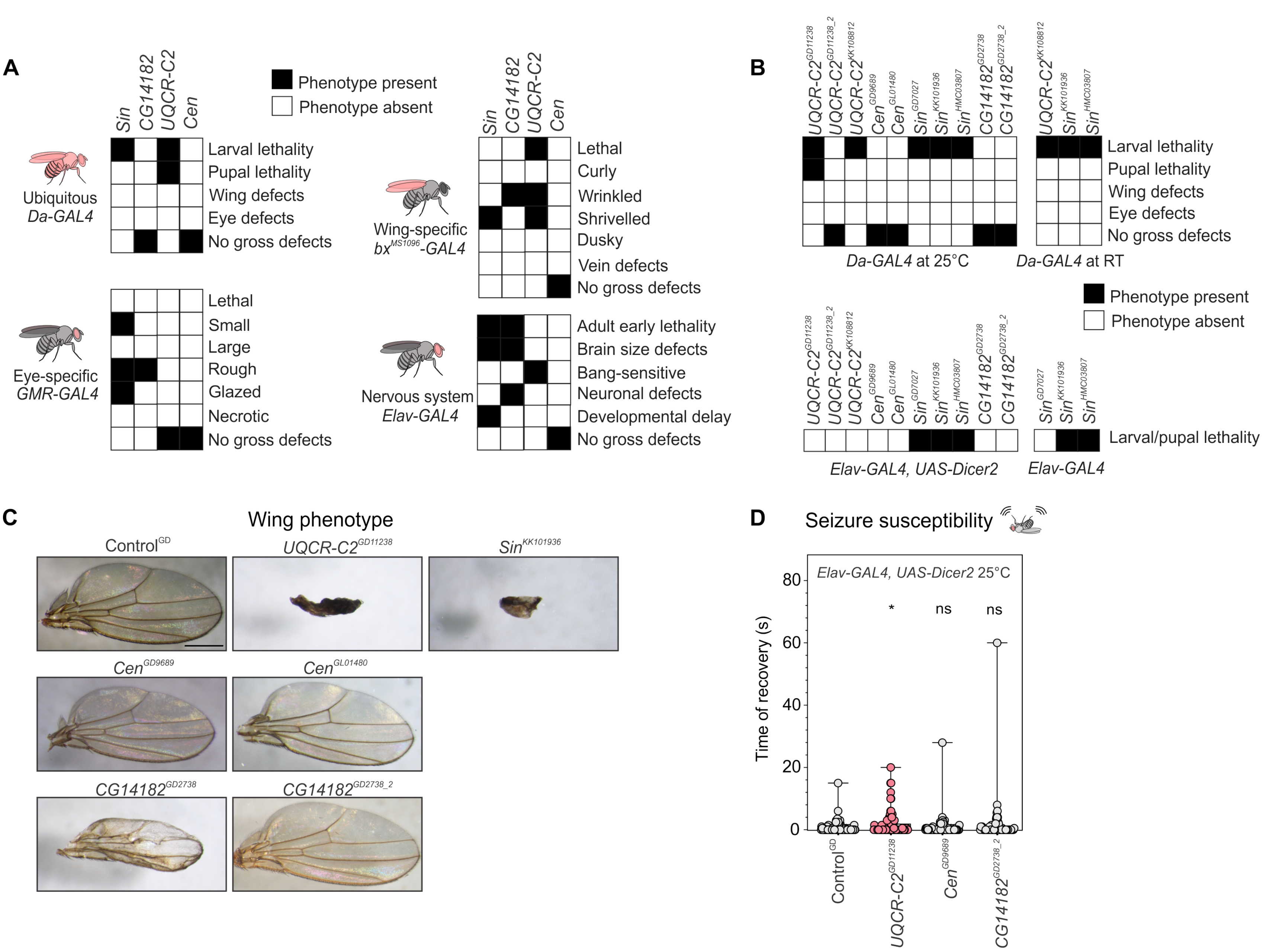

### S3 Figure.pdf

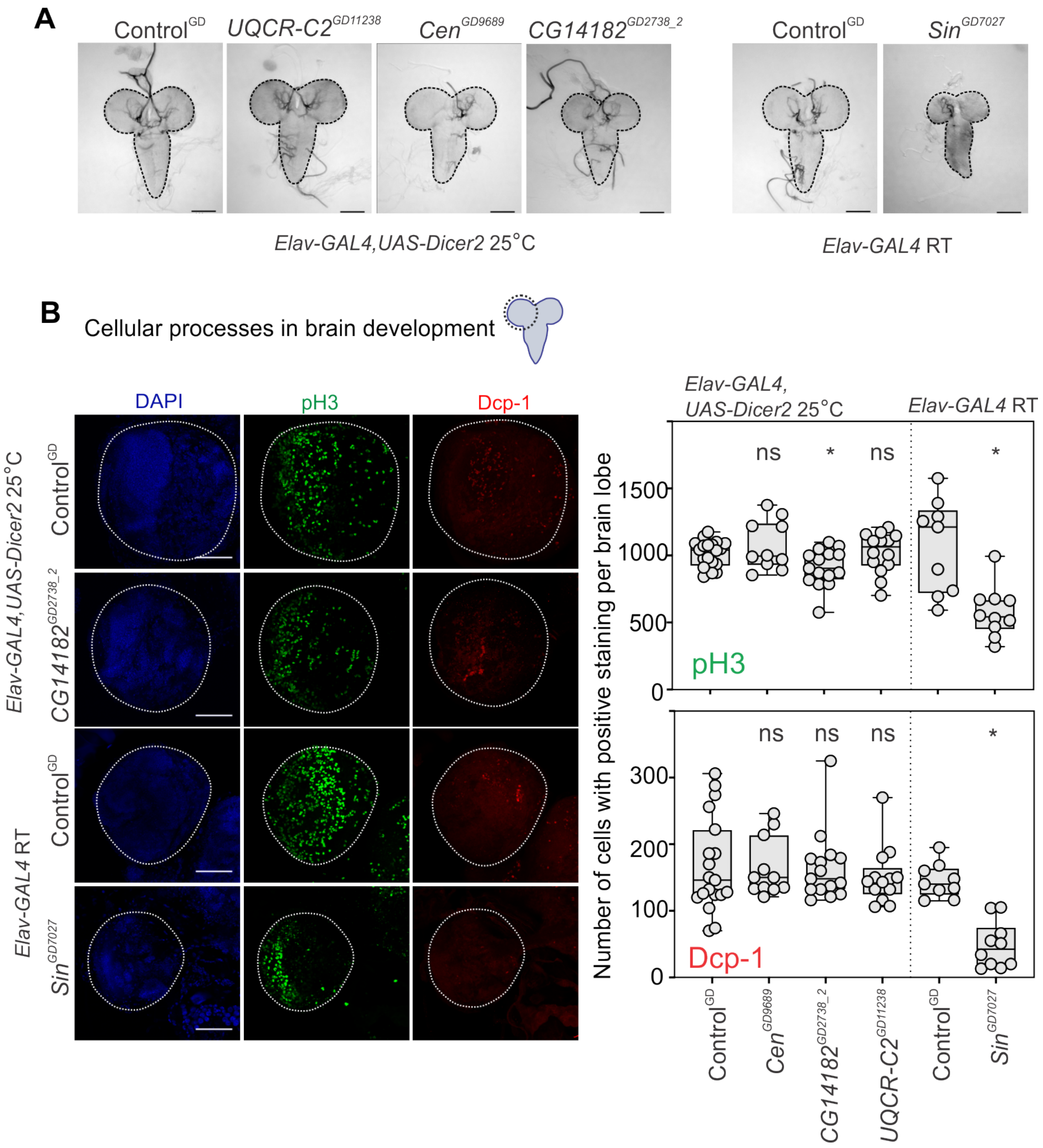

### S5 Figure.pdf

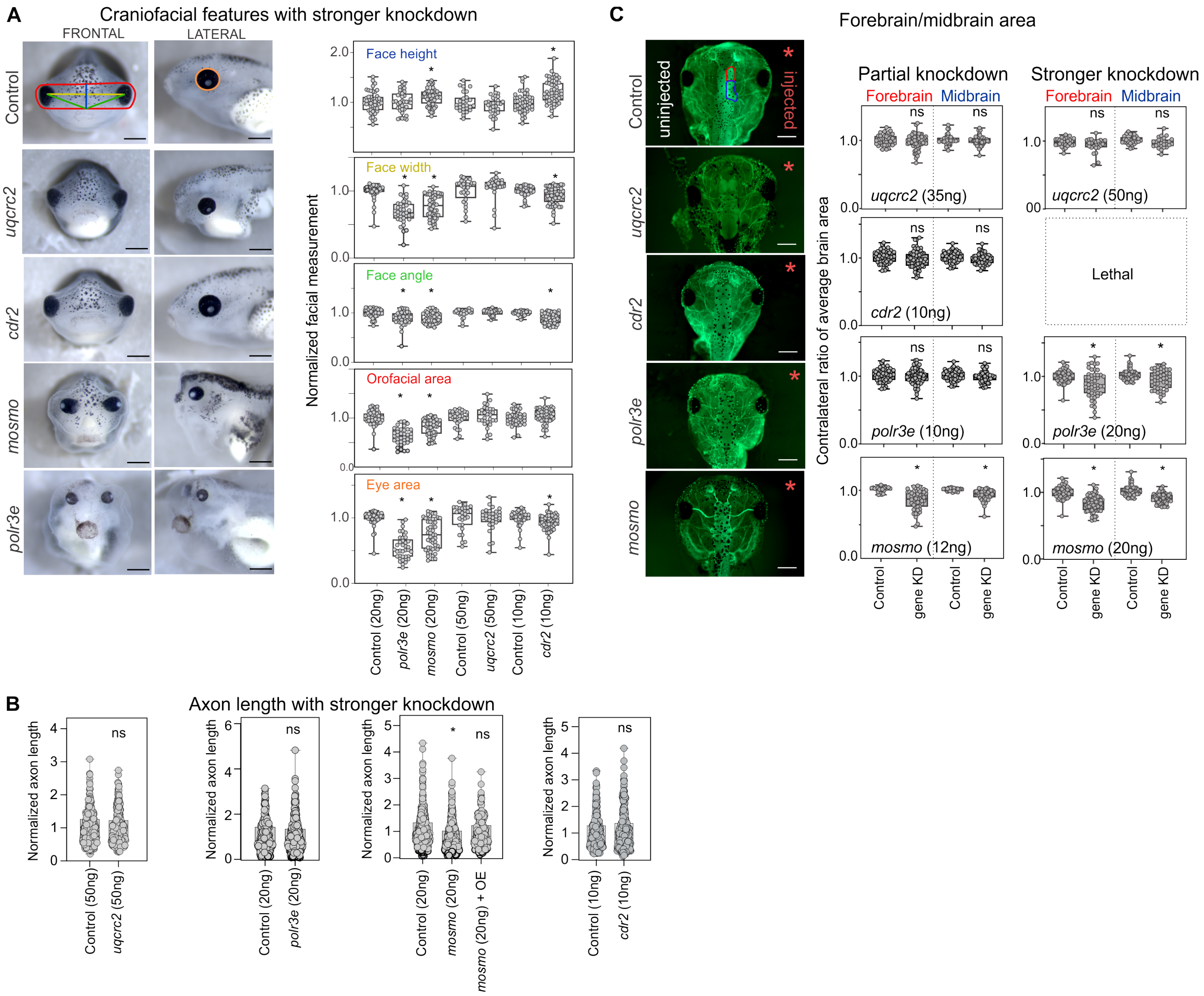

### S6 Figure.pdf

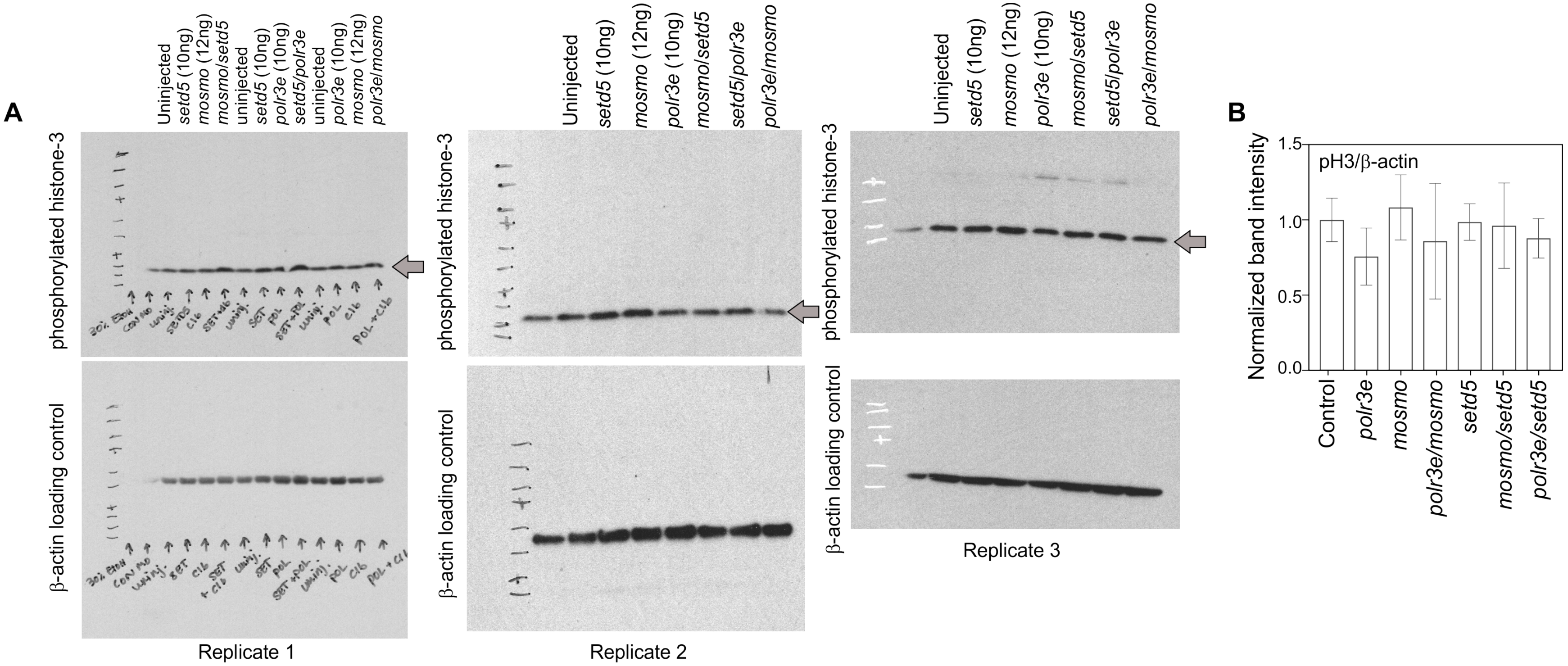

### S7 Figure.pdf

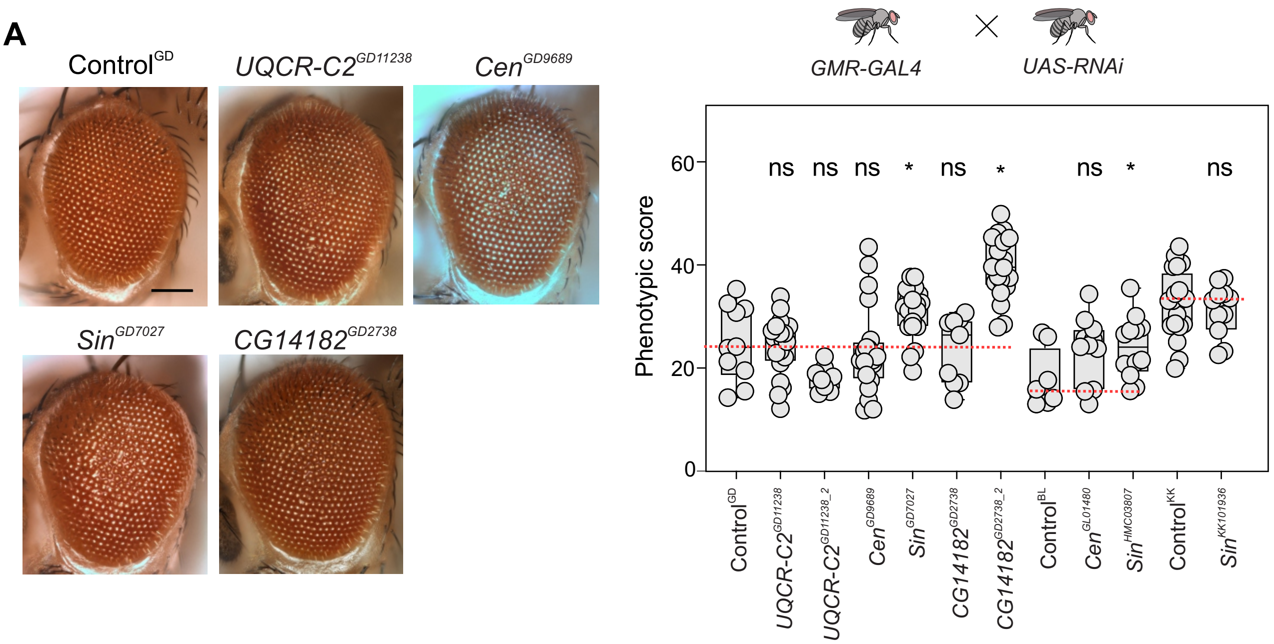

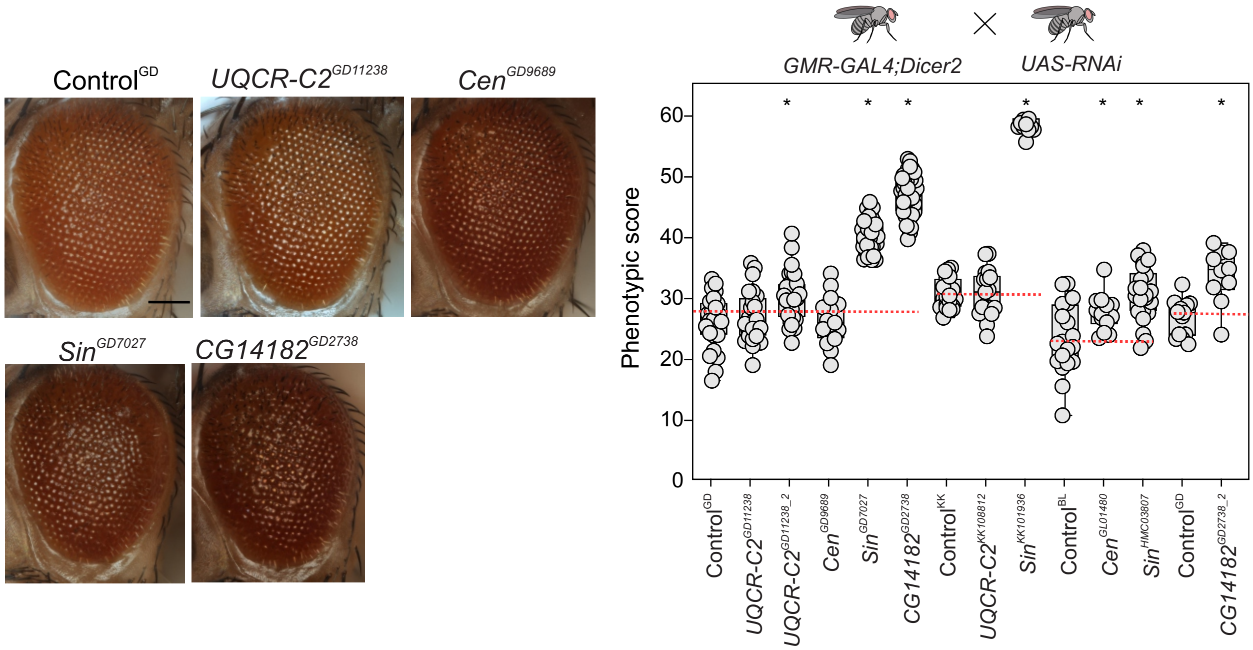

### S8 FIgure.pdf

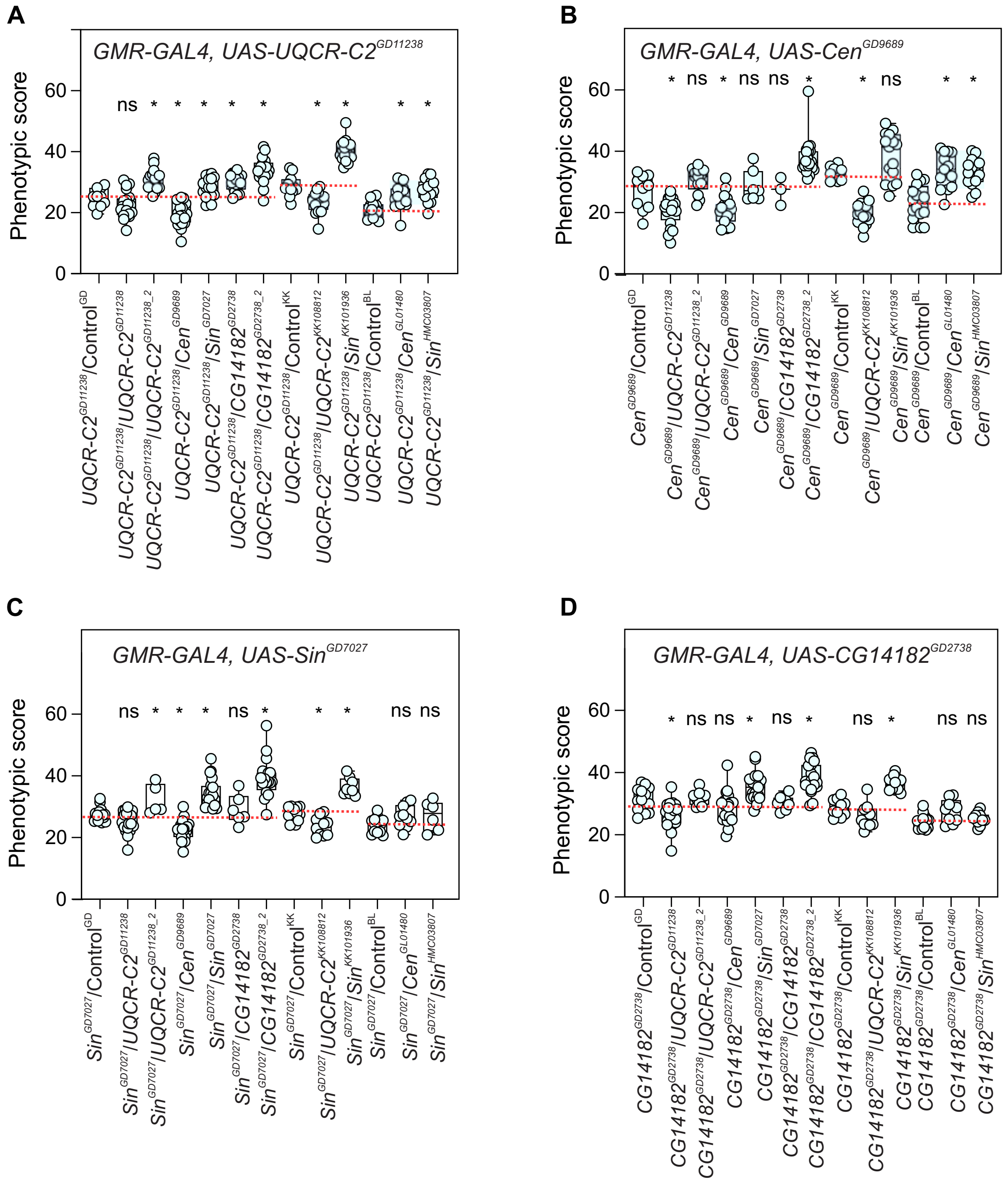

### S9 Figure_revision.pdf

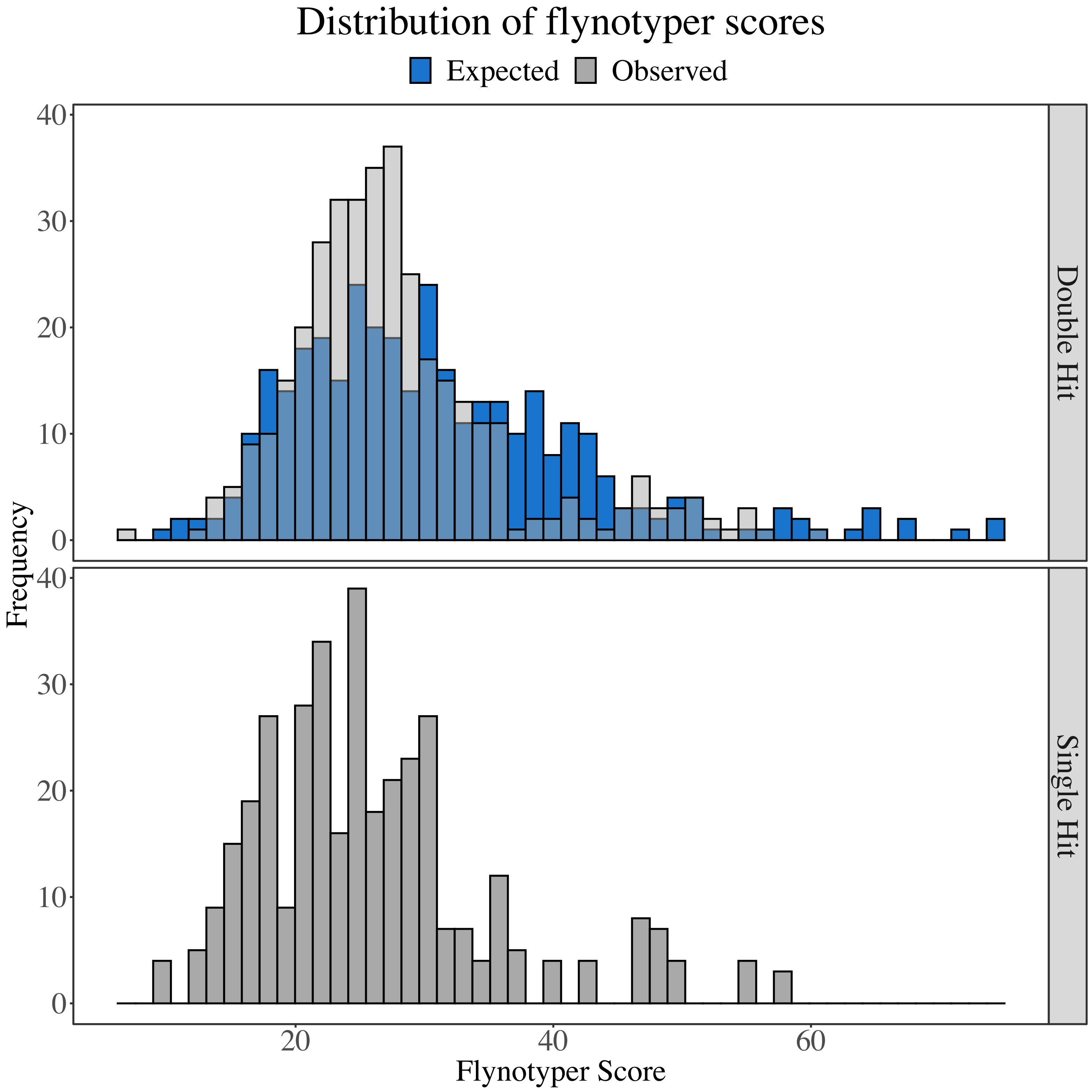

### S10 Figure_revision.pdf

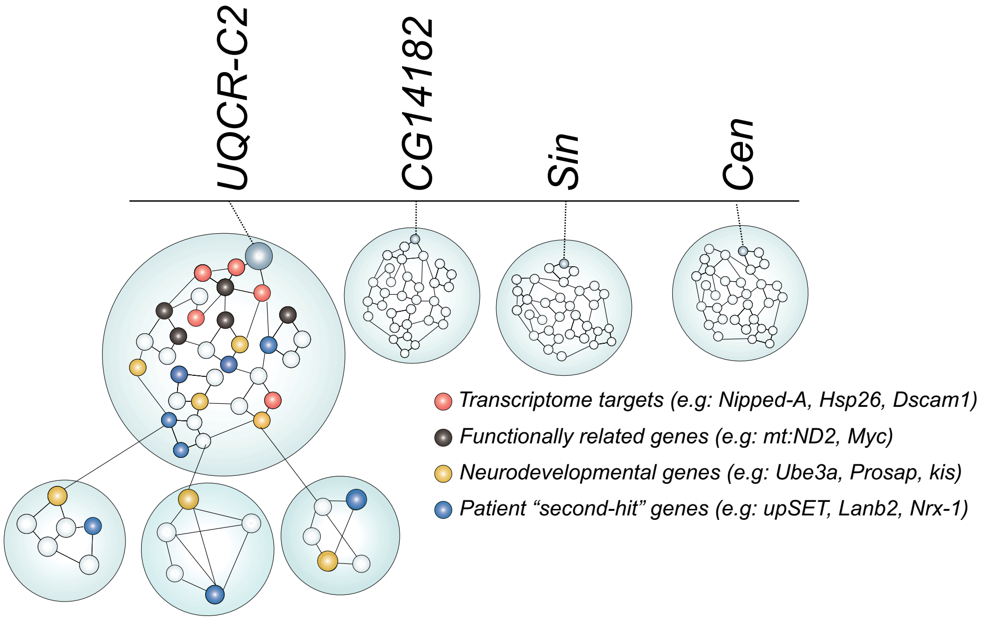

### S11 Figure_revision.pdf

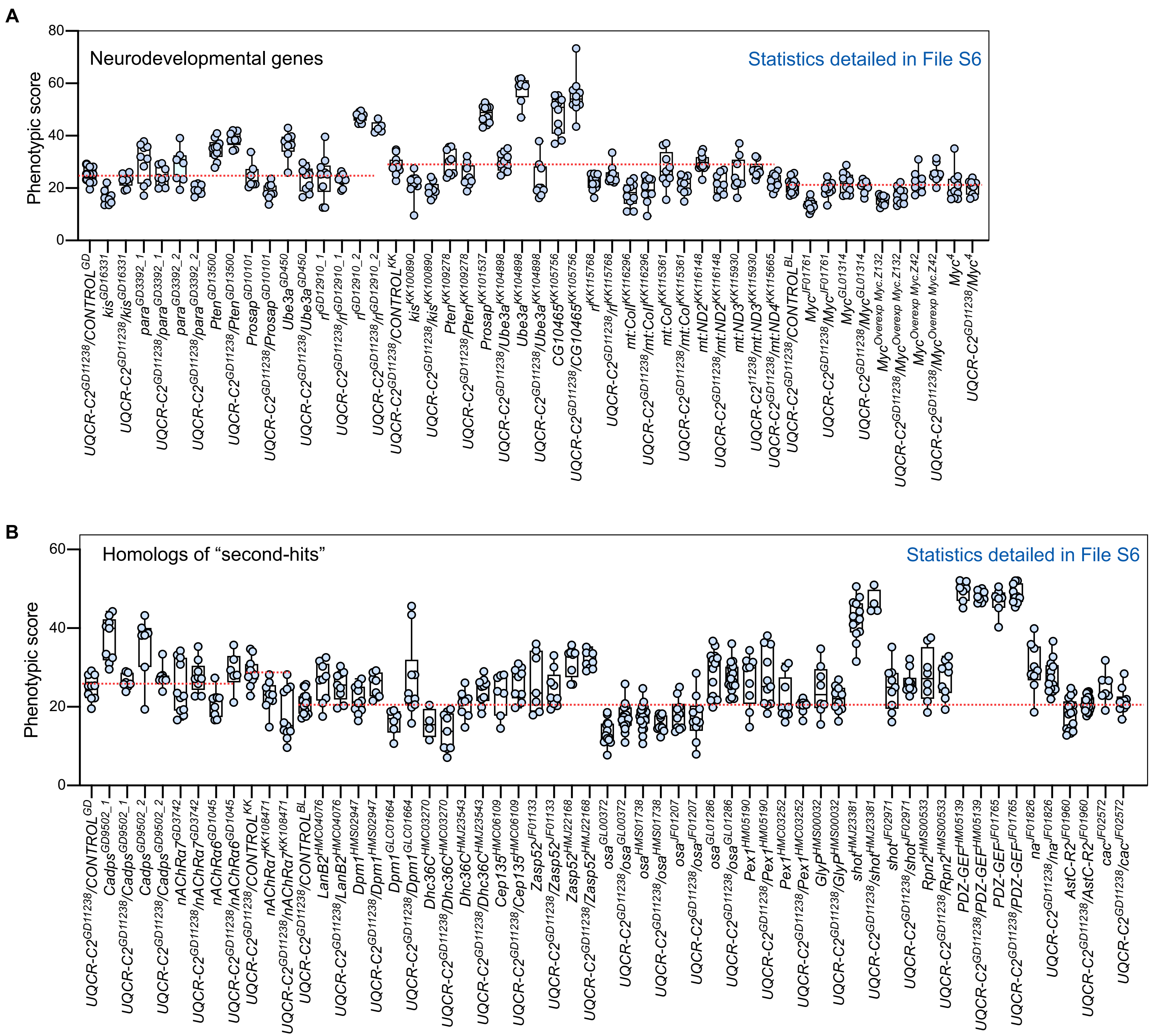

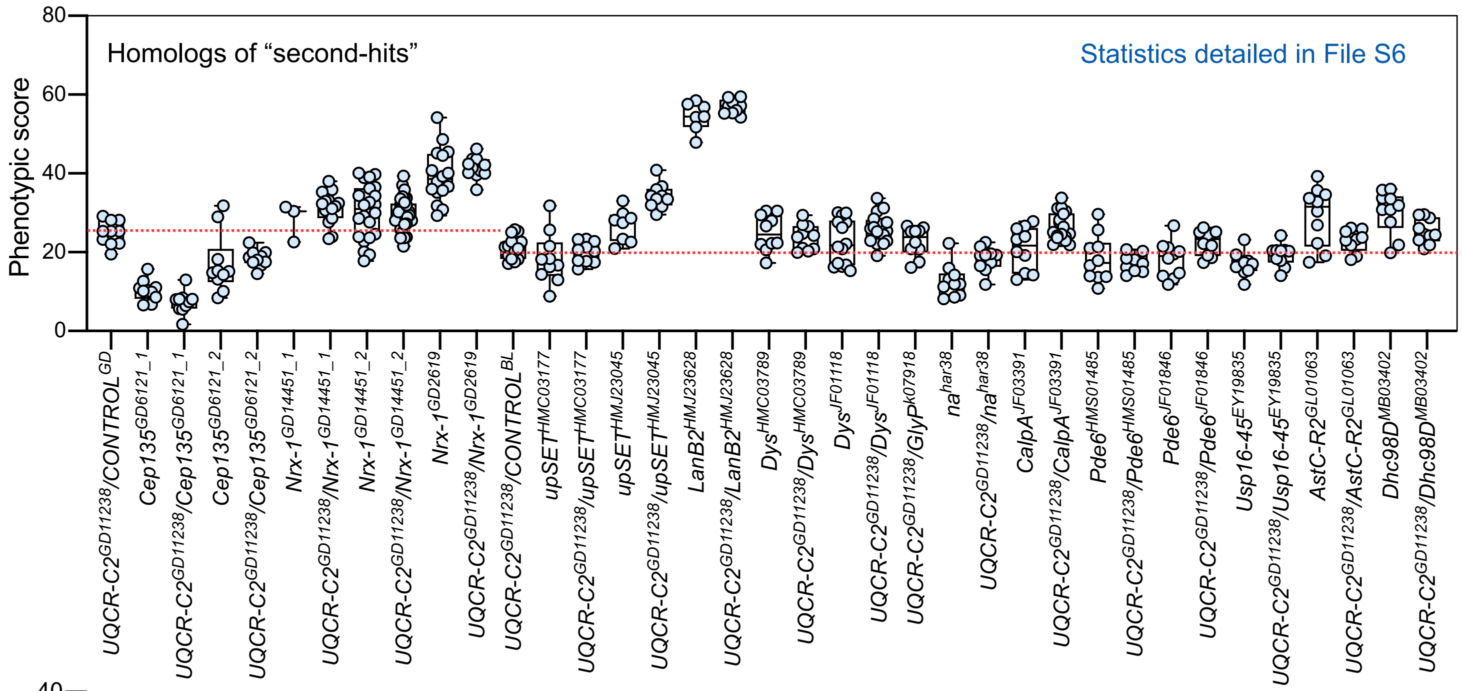

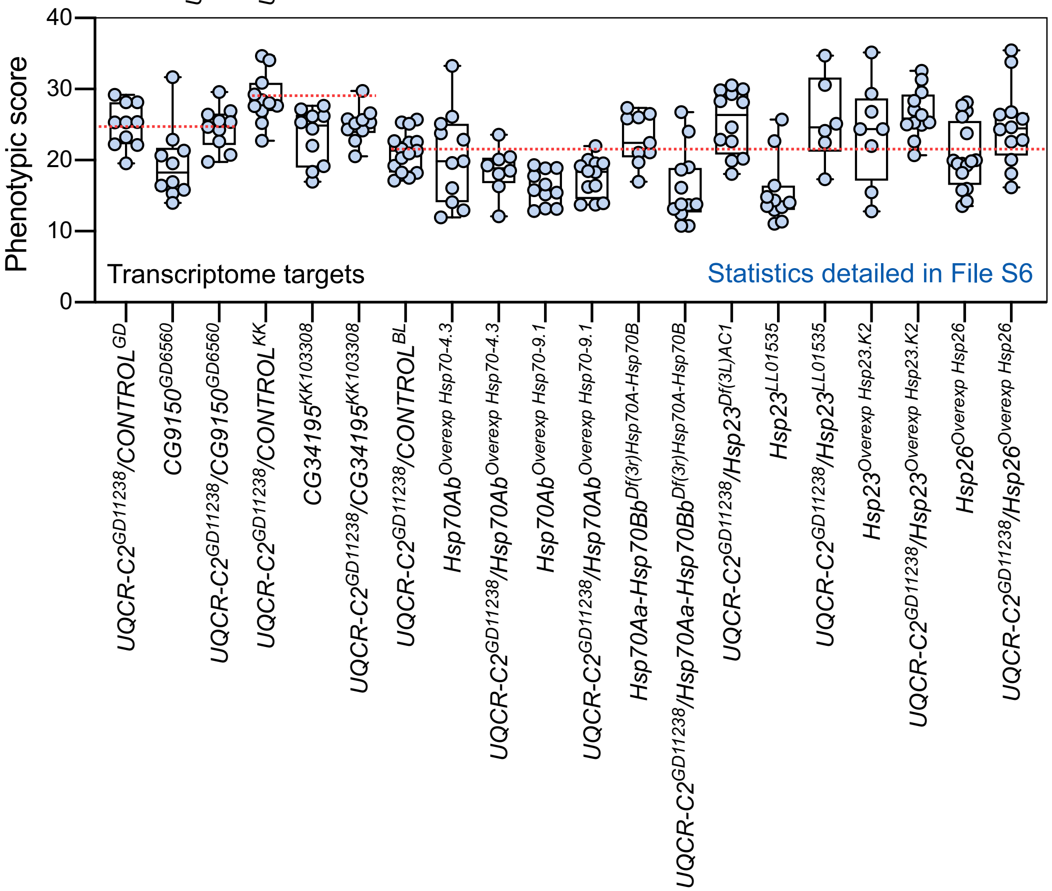

### S12 Figure_revision.pdf

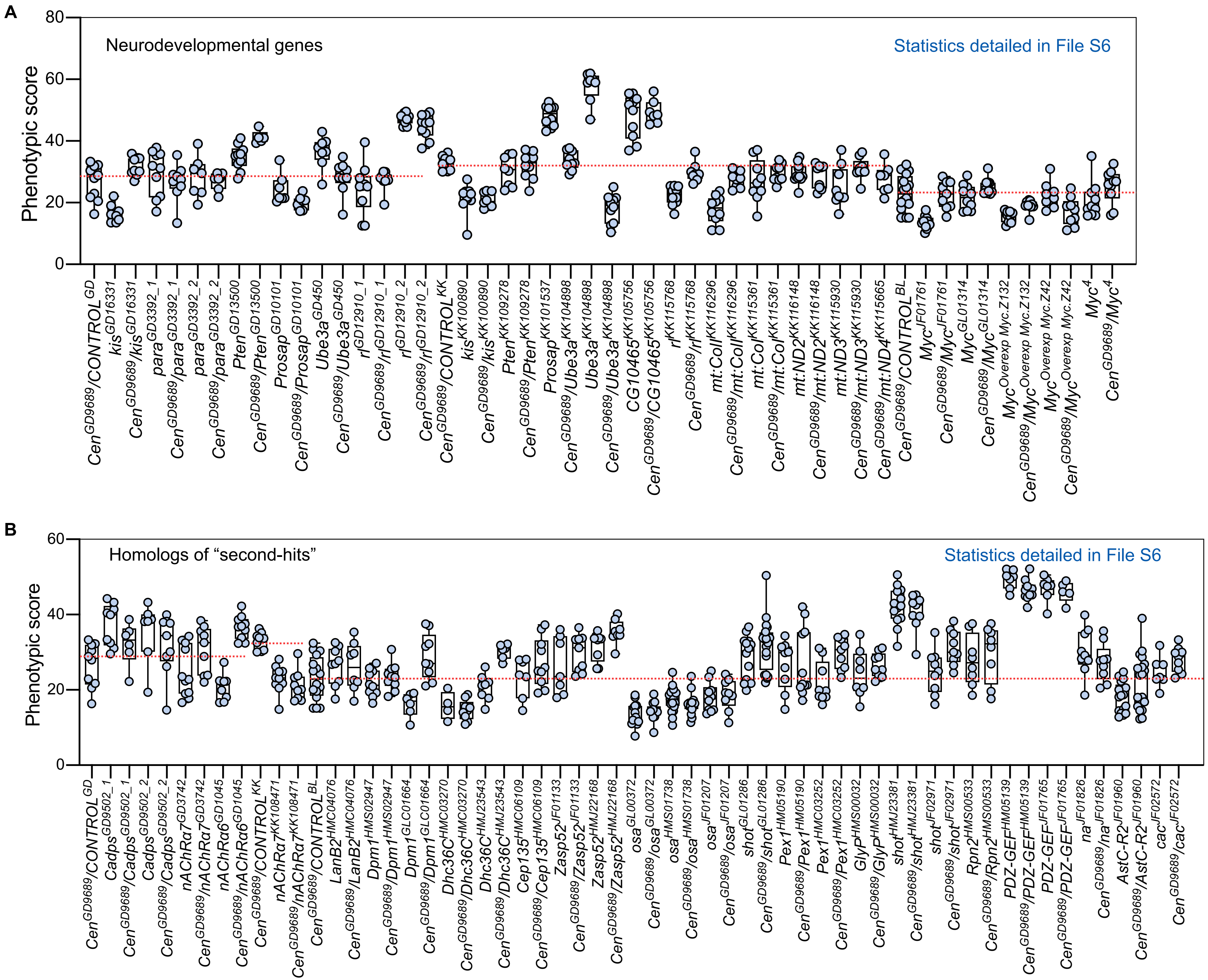

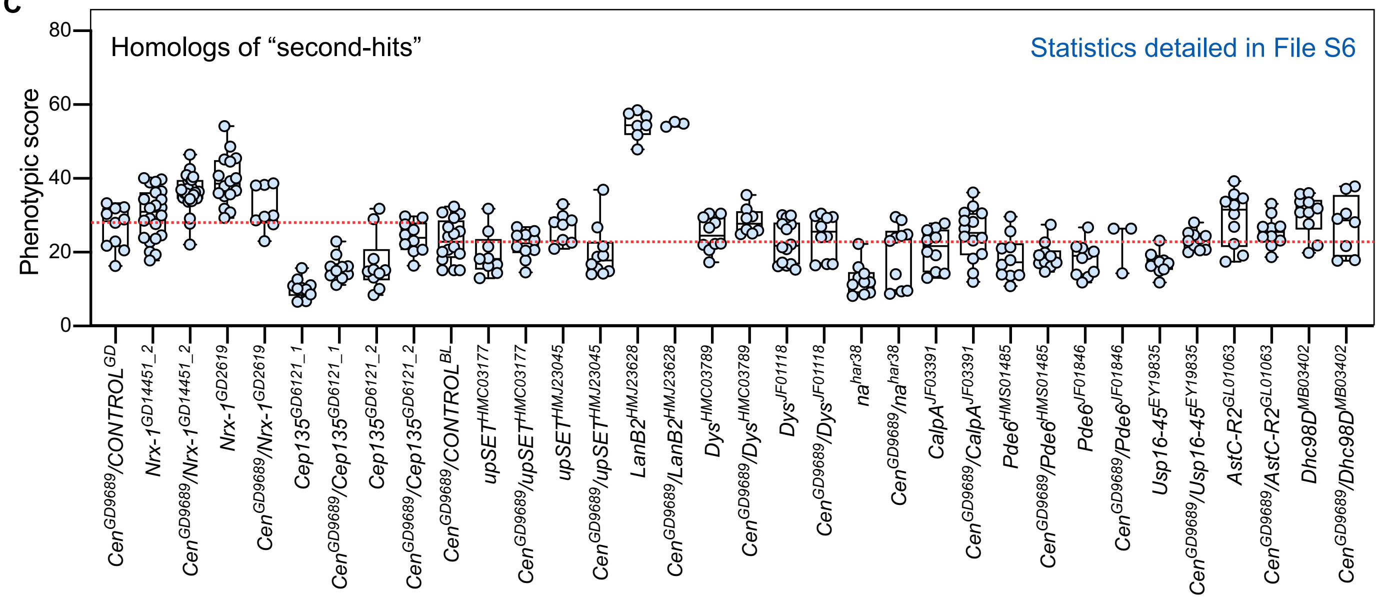

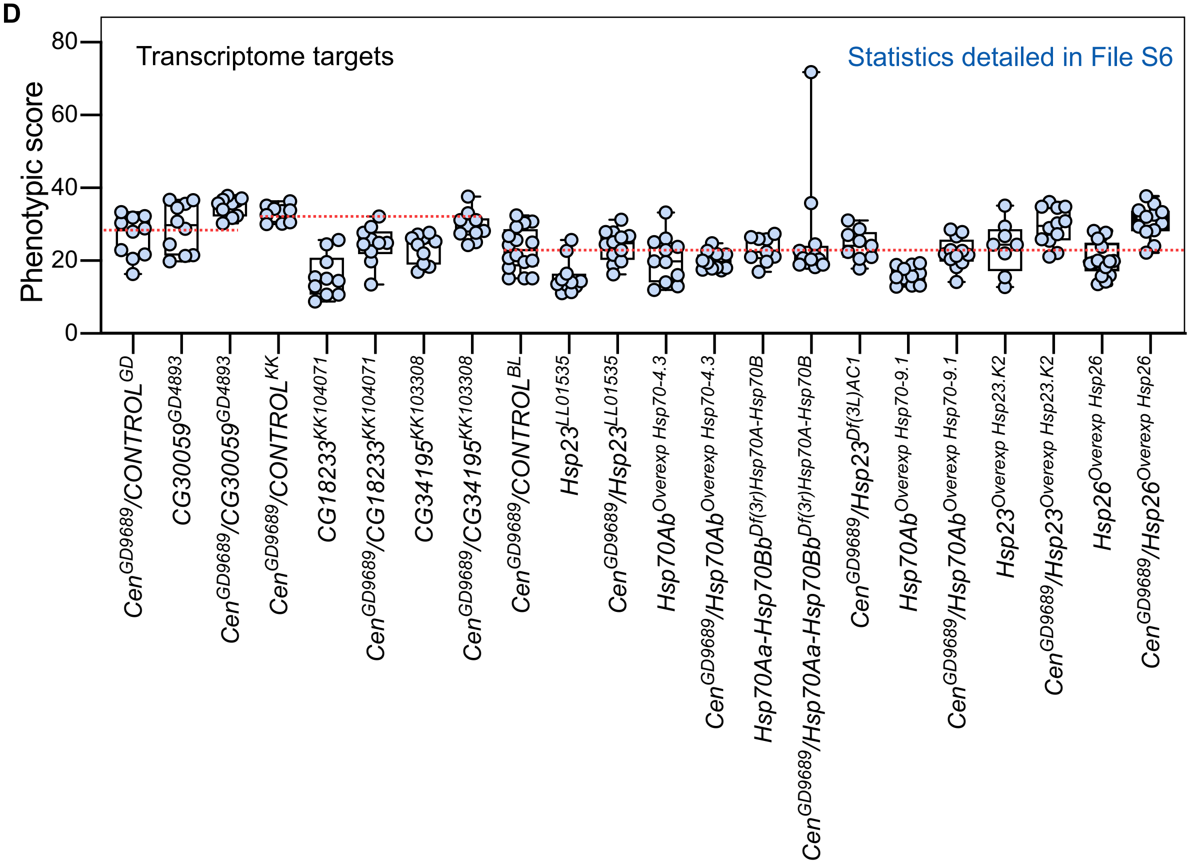

### S13 Figure_revision.pdf

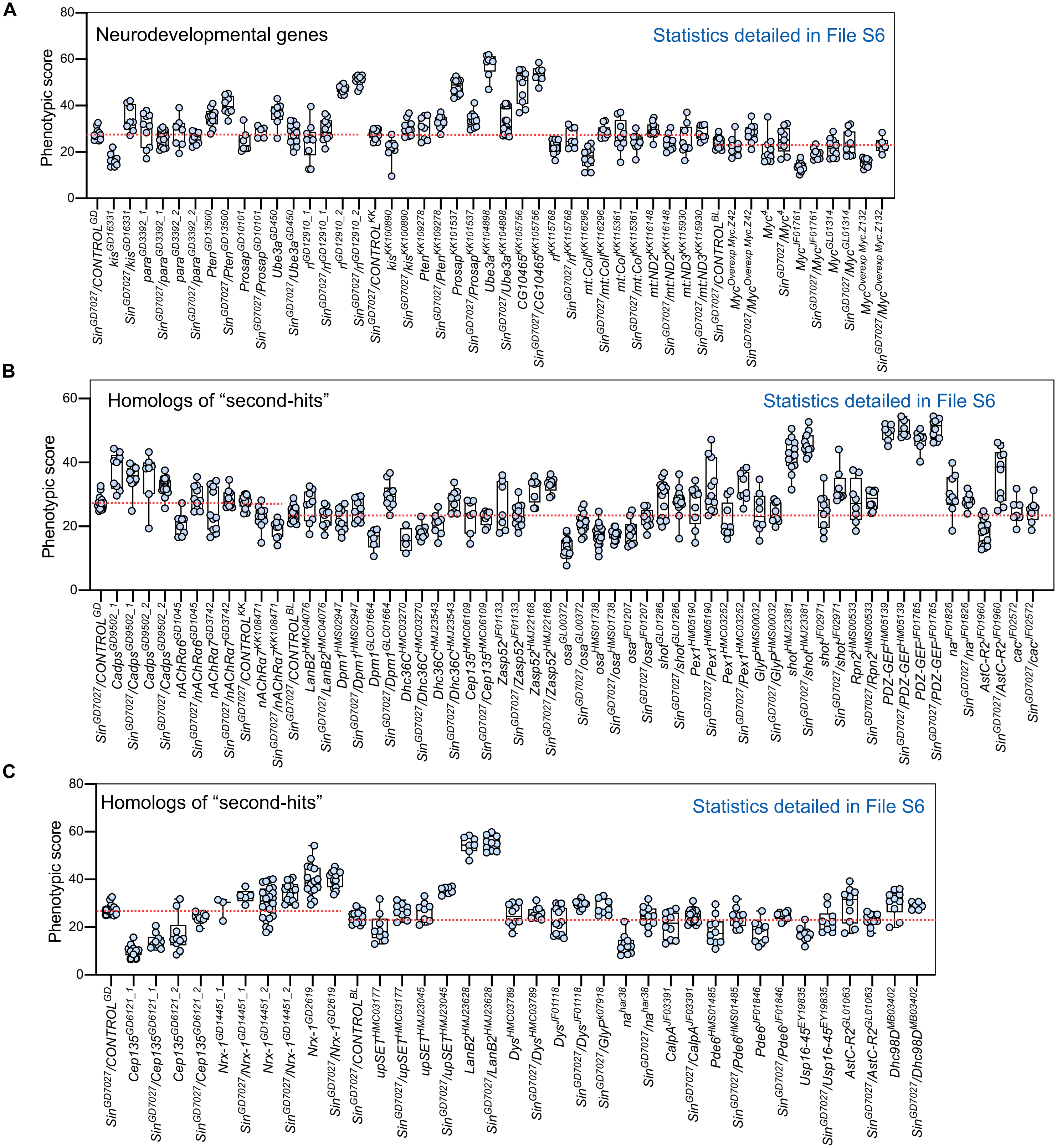

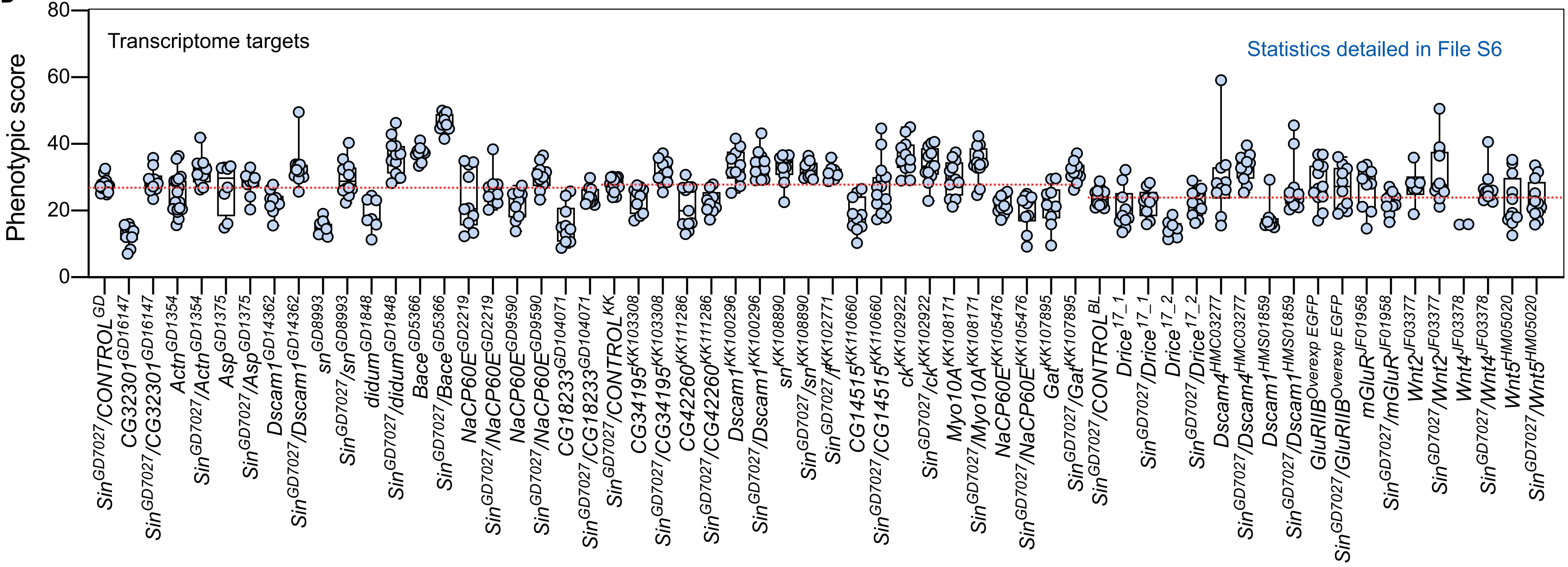

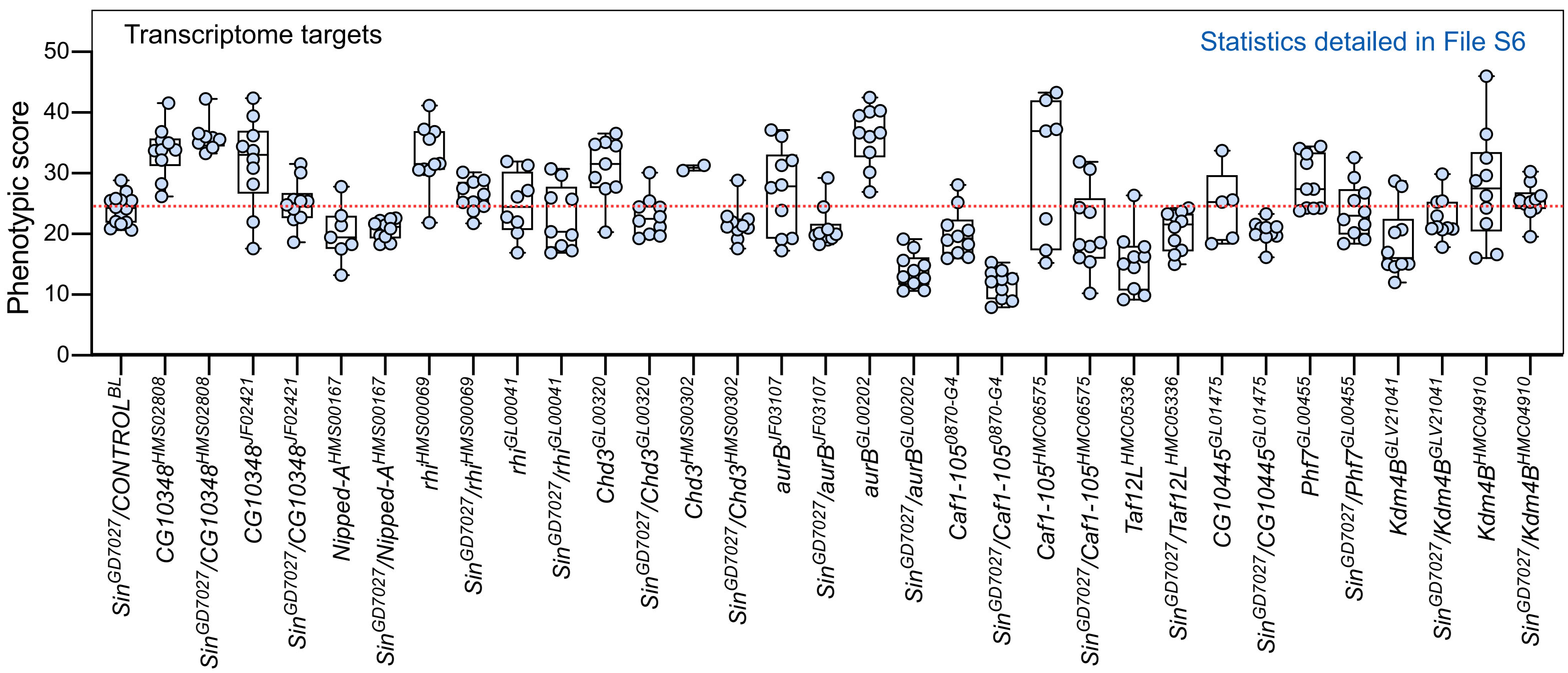

### S14 Figure_revision.pdf

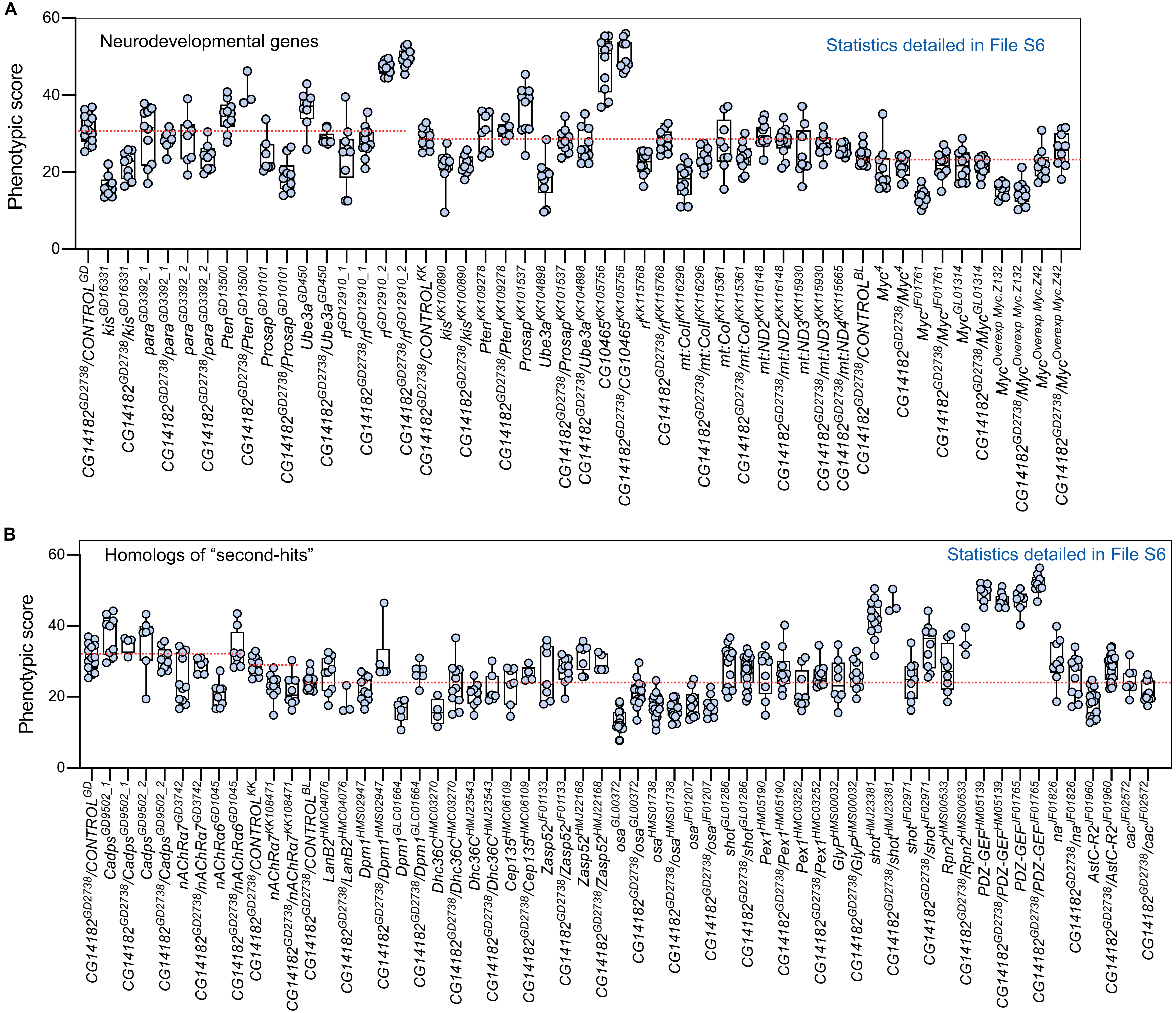

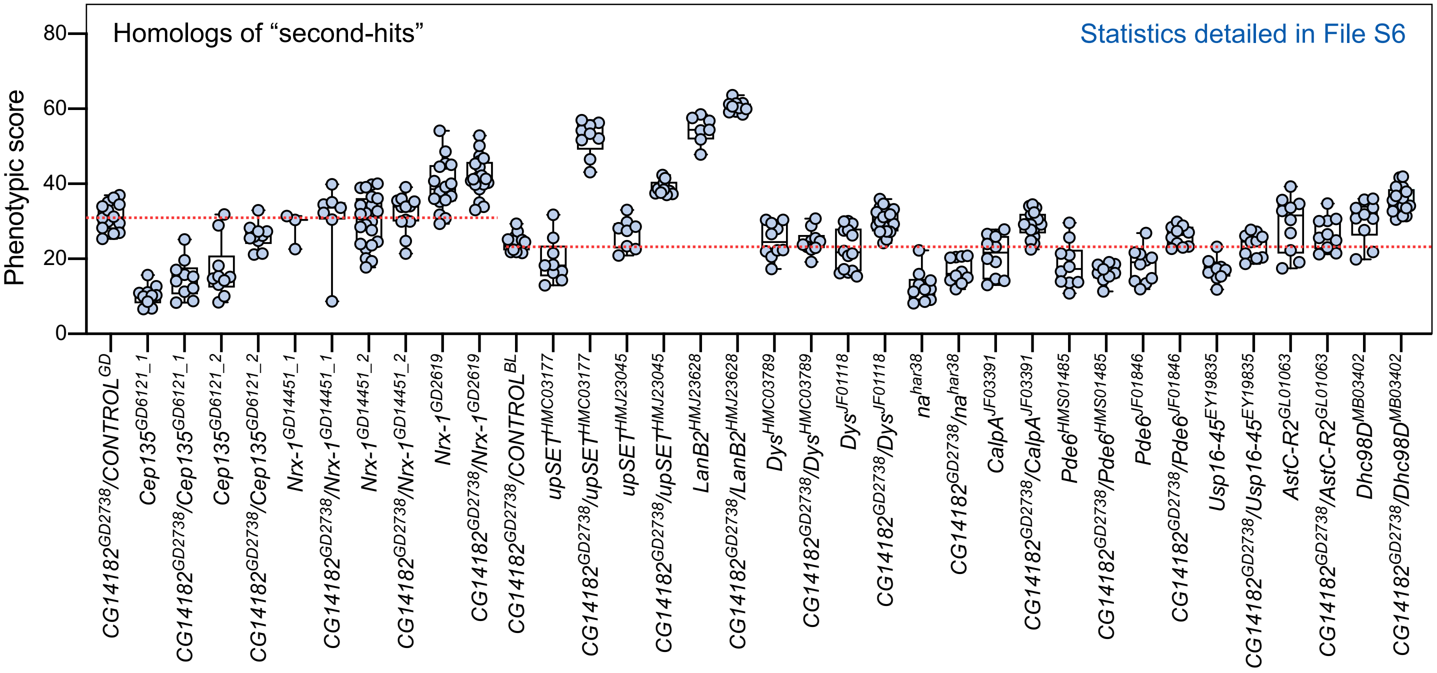

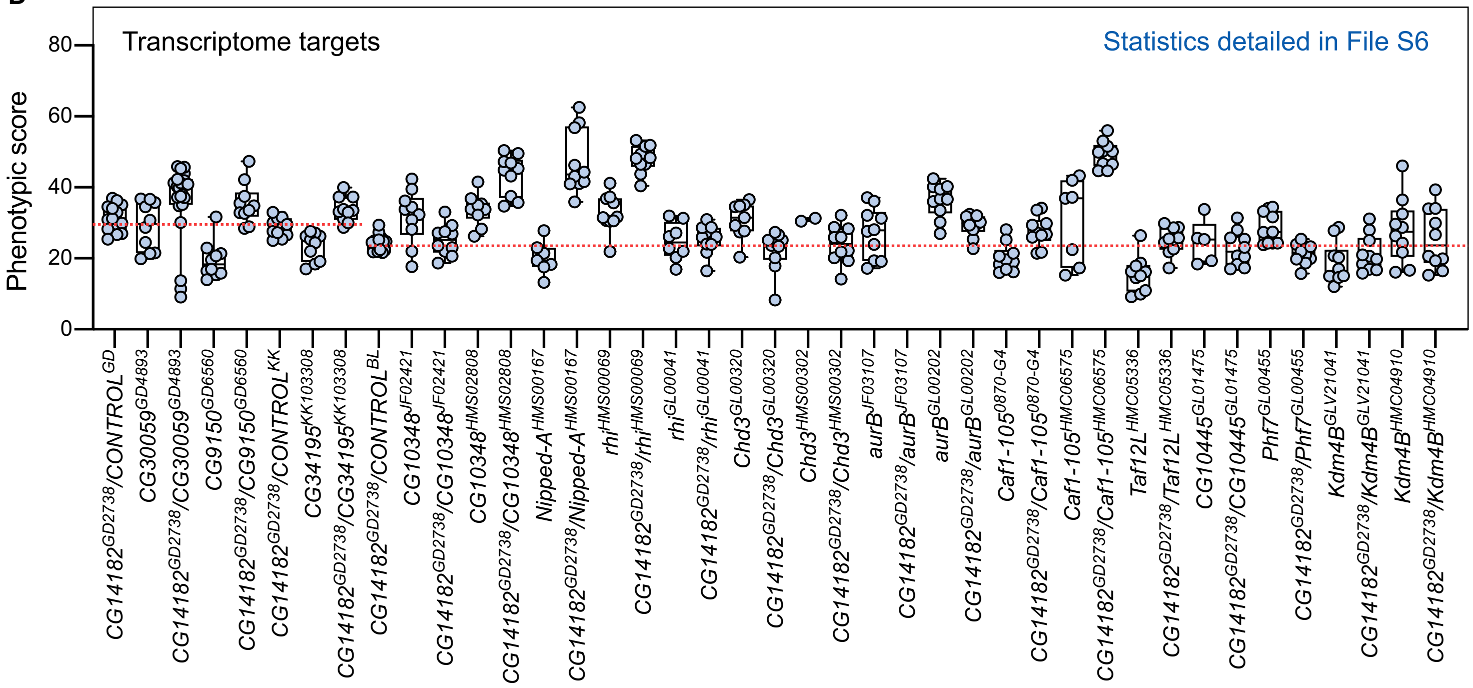

### S15 Figure_revision.pdf

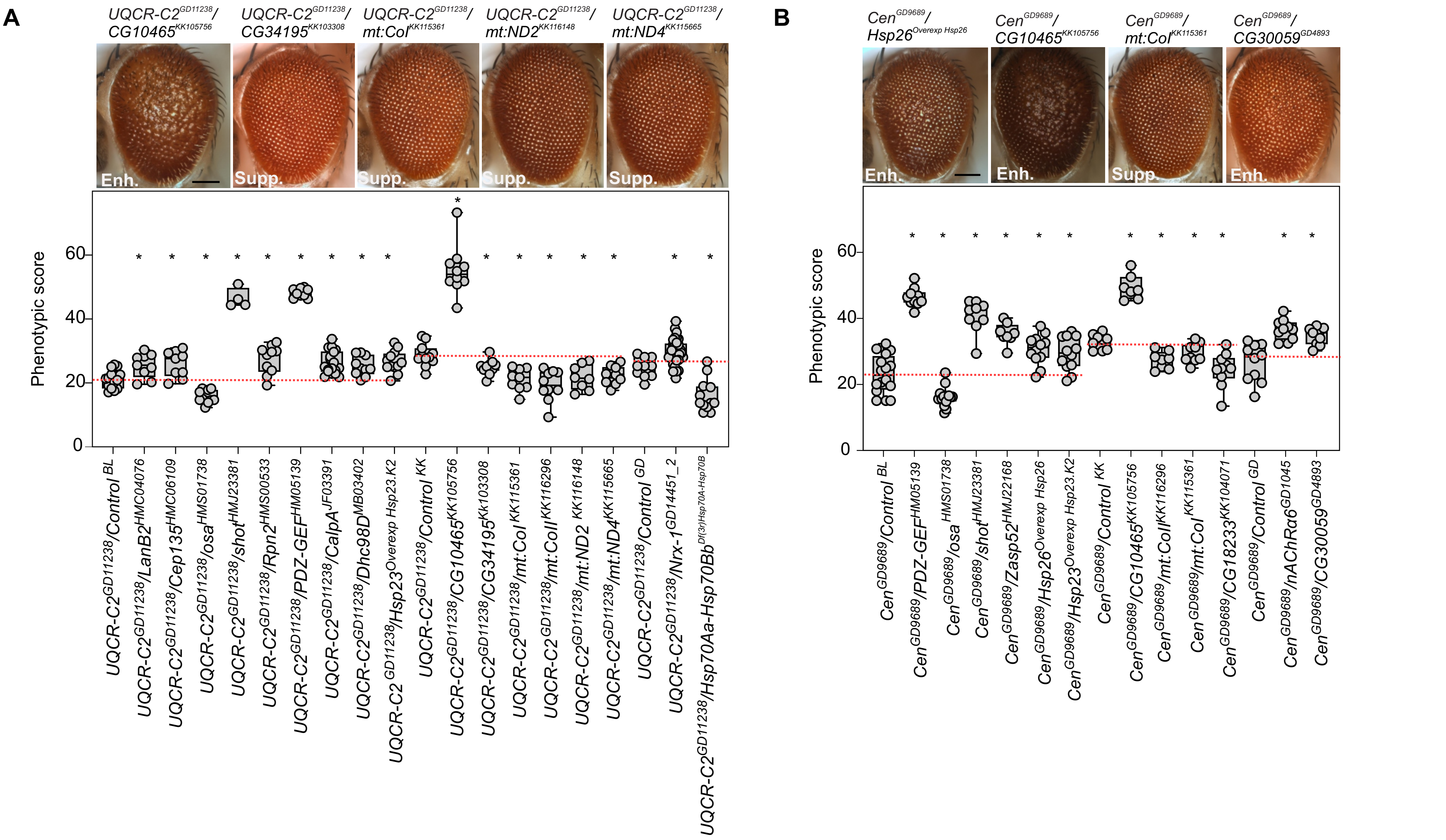

### S16 Figure_revision.pdf

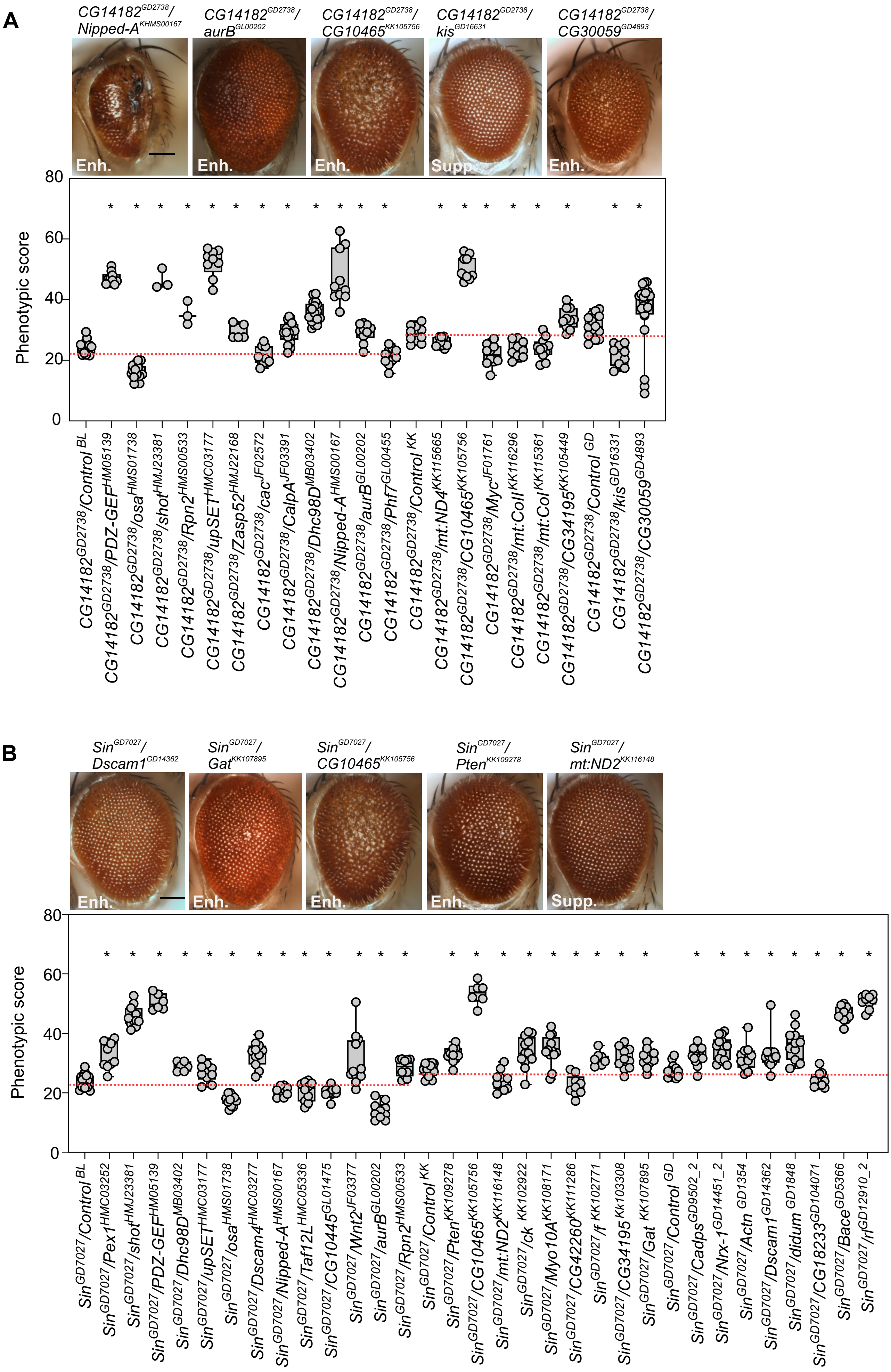

### S17 Figure_revision.pdf

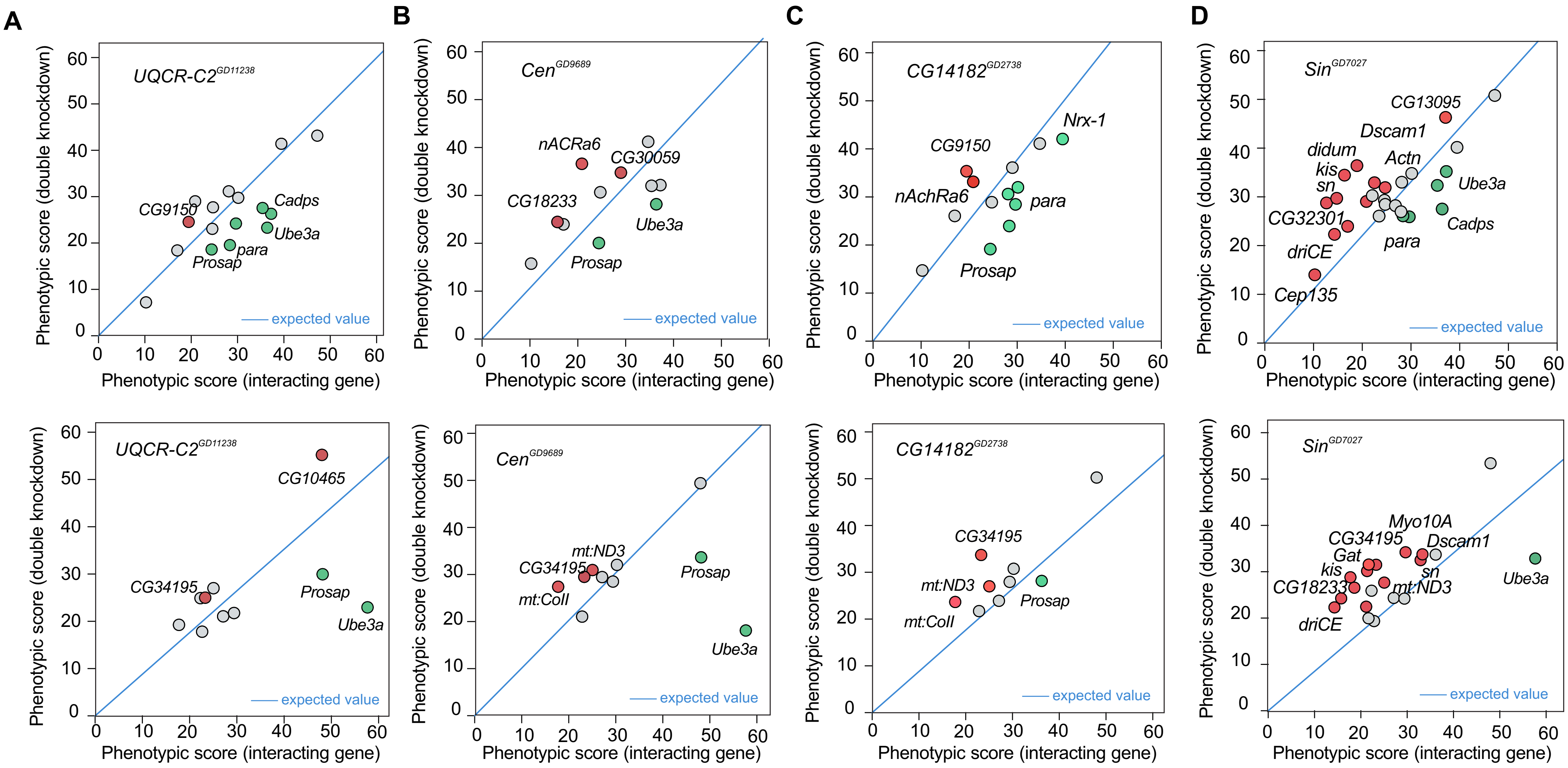

### S18 Figure_revision.pdf

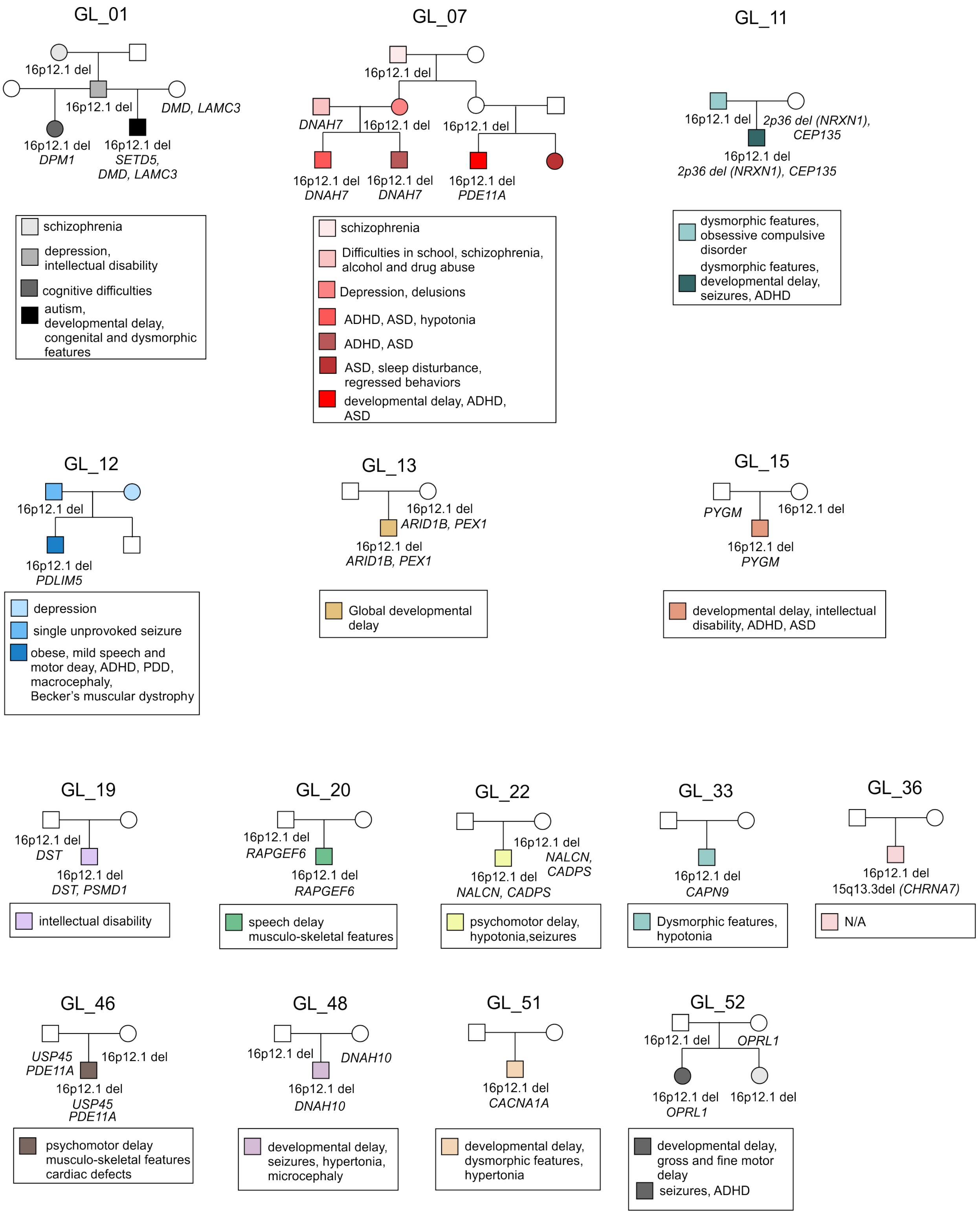

### S19 Figure_revision.pdf

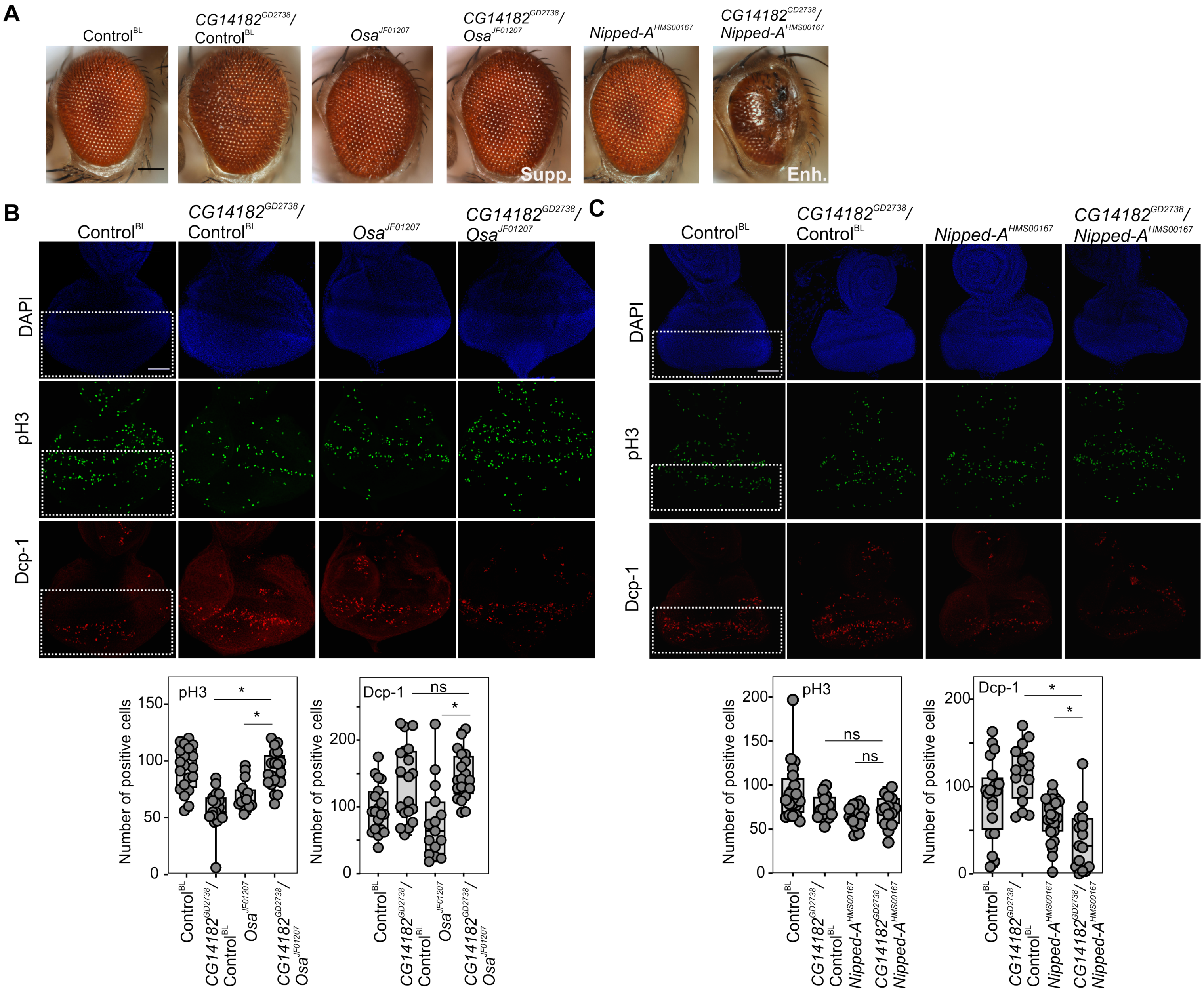

### S20 Figure_revision.pdf

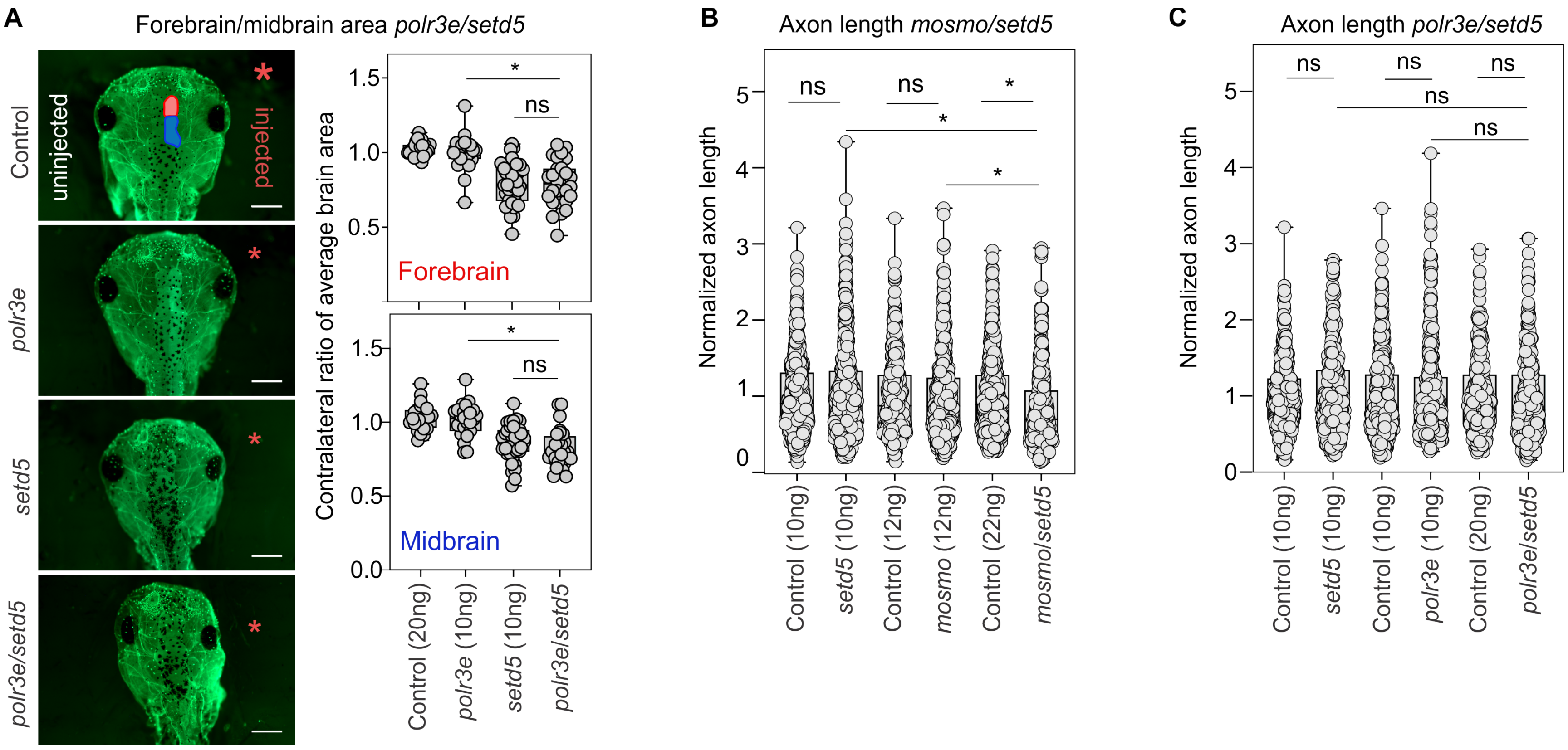

### S21 Figure_revision2.pdf

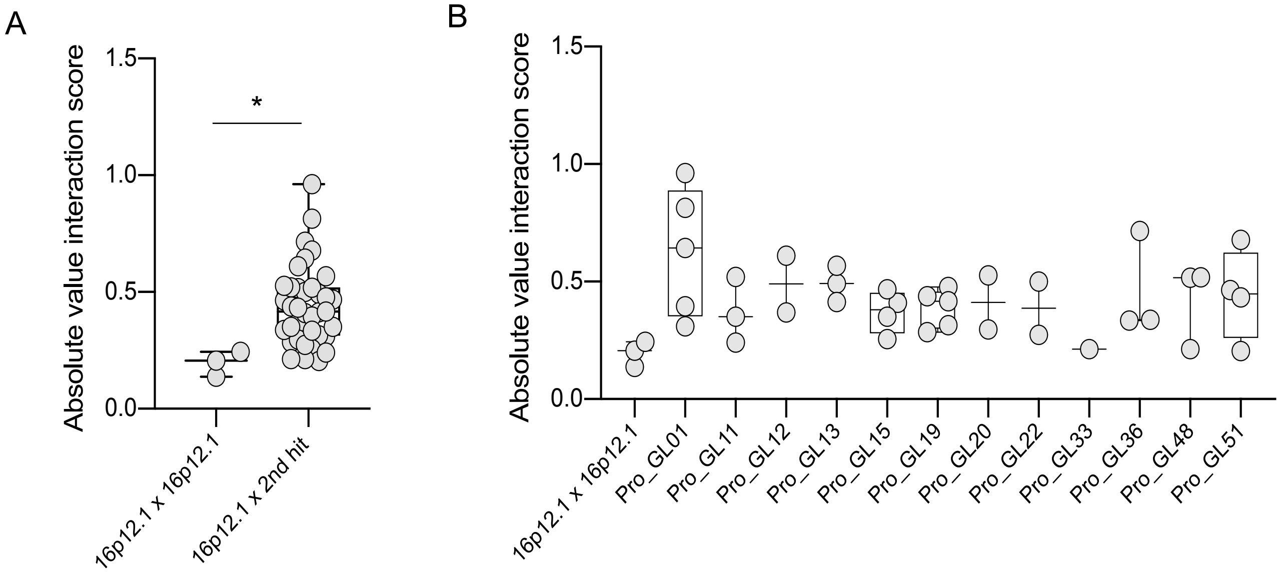

### S22 Figure_revision2.pdf

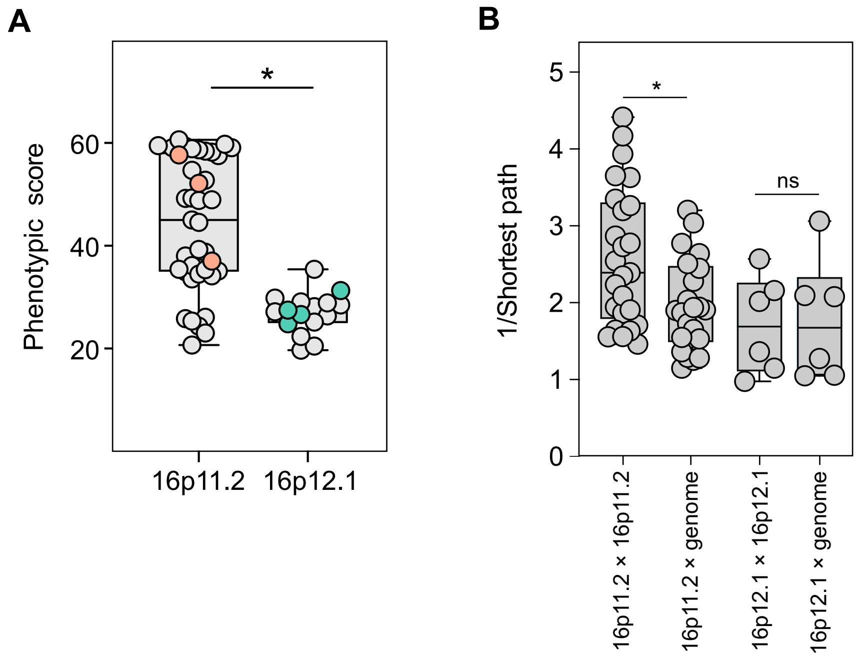

### S23 Figure_revision2.pdf

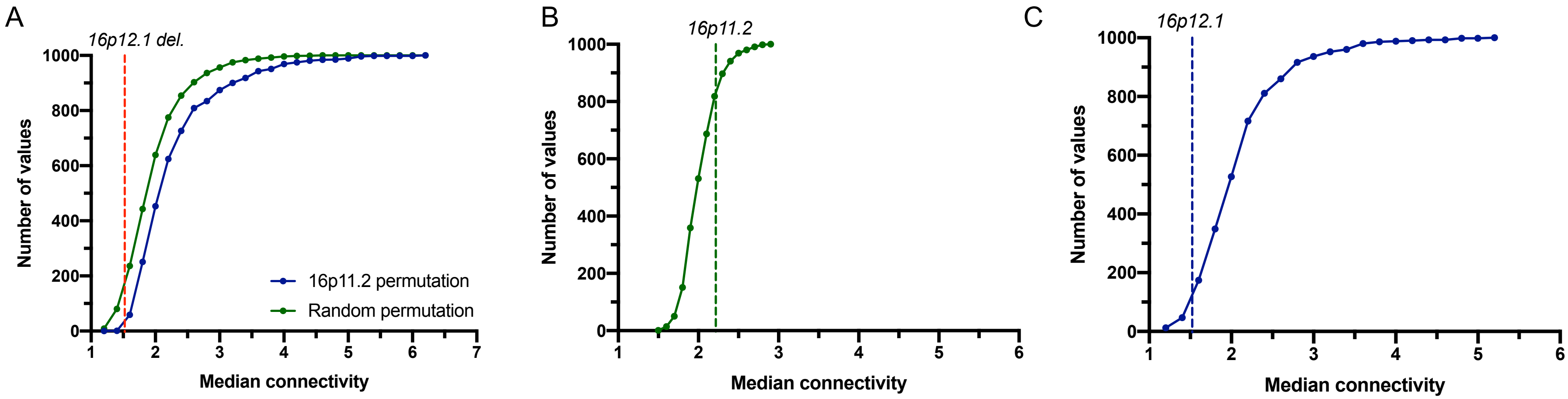
